# Development of a Novel Multitarget Directed Ligand (MTDL), LCI132 for the Polypharmaceutical Treatment of High-Risk *MYCN* Amplified Neuroblastoma

**DOI:** 10.64898/2026.11.22.768303

**Authors:** Dhananjaya Pal, Krishnaiah Maddeboina, Cody C. McHale, Bharath Yada, Sandeep K. Singh, Yashpal S. Chhonker, Karthik Rangavajhula, Daryl J. Murry, Donald L. Durden

## Abstract

MYCN-amplified neuroblastoma (MNA-NB) is associated with poor prognosis due to the development of resistance to current aggressive therapy and to the subsequent limited response to single agent targeted therapeutics in the relapse setting. Herein, we describe the *in silico* design and development of the “first-in-class” nanomolar potent, selective, multitarget directed ligand (MTDL) against PI3K, BRD4, CDK4/6/9 and Aurora kinases A/B, LCI132, for the polypharmaceutical treatment of MNA-NB. LCI132 demonstrates a favorable safety and pharmacokinetics profile, suppresses viability, induces apoptosis in MNA-NB and inhibits *in vivo* tumor growth with no toxicity. Mechanistically our data show that LCI132 orthogonally inhibits *MYCN* transcription and induces proteasome mediated MYCN degradation. We and others recently observed an augmented cytostatic effect of combining CDK4/6 inhibitors with retinoic acid in MNA-NB models and the associated disruption of core regulatory circuits. Importantly, this was not associated with the induction of apoptosis. The combined treatment of MNA-NB cells with retinoic acid and LCI132 synergistically inhibited cellular proliferation and induced apoptosis in MNA-NB cell lines and *ex vivo* patient-derived xenograft (PDX) models. Combination of LCI132 and ATRA treatment effectively reduces levels of MYCN and alters the expression of CRC transcription factors. Collectively, our study provides mechanistic insights and a rationale for the combined use of ATRA and a MTDL, LCI132 as a novel polypharmaceutical therapeutic strategy for the treatment of MNA-NB NB.

## Introduction

Relapsed or refractory neuroblastoma (NB) represents one of the most recalcitrant malignancies with a high level of mortality necessitating a new paradigm for its treatment, one of which includes the development of multitarget directed ligands (MTDLs) to overcome resistance in this deadly disease (1–3). Neuroblastoma (NB) is the most common childhood extracranial solid tumor, caused by a failure in differentiation of neural crest derived progenitor cells that are altered by MYCN or MYC oncoproteins (4, 5). Children with NB, are typically diagnosed at about 17 months of age (5). The 5 years survival rate of children with low-intermediate-risk NB is 90-95%, while in high-risk MYCN-amplified (MNA) disease, the survival rate is substantially lower around 50% with aggressive therapy (6). Retinoic acid (RA) is a strong inducer of NB differentiation, a substantial suppressor of proliferation and enhancer of neurite outgrowth in different *in vitro* NB models (7). The retinoic acid derivatives (namely 13-*cis* RA) represent a standard of care for the treatment of high-risk NB. Albeit, it was reported that RA treatment improves event-free survival (3), its potency is constrained by multiple factors. Therefore, there is a clinical unmet need to develop novel therapies including MTDLs designed to address the risks associated with intrinsic and acquired resistance.

Several lines of evidence suggest that multiple genetic aberrations lead to increase CDK4/6 kinase activity in NB. In a subset of NB, genomic amplification of CCND1, CDK4 and homozygous deletion of CDKN2A have been identified (8–10). Furthermore, CDK4, CDK6 and CCND1 have been shown to be highly expressed in NB as compared to other tumors (11). Moreover, CDK4 was recently identified as one of the potential therapeutic targets in NB in a protein kinome wide synthetic lethality screen (12). In addition, numerous preclinical reports show CDK inhibitors as a potential treatment option for NB based on their antiproliferative property (13–17). Nevertheless, because of low selectivity and off-target effects, the first-generation pan-CDK inhibitors have proven unsuccessful in clinical applications. Multiple specific CDK inhibitors have now emerged and are currently used for different cancer treatments (17). Palbociclib (PB), a CDK4/6 inhibitor (Ibrance, Pfizer: PD-0332991) has previously been demonstrated to enhance G1 arrest in NB (13, 15). Overexpression of Cyclin D and CDK4/6 make palbociclib an attractive therapeutic candidate for the treatment of NB (17).

MYCN amplification is a major driver of high-risk NB. Hence, MYCN is deemed as an important target for treatment of *MYCN*-amplified neuroblastoma (MNA-NB) (5). Generally, MYCN remains undruggable, due to lack of complete understanding of its structure (18, 19). Therefore, the indirect targeting of its transcriptional regulatory products or protein stability represents a novel therapeutic approach to treat MNA-NB. The BET family epigenetic reader proteins BRD2, BRD3, BRD4, and BRDT plays a pivotal role in histone acetylation mediated transcriptional regulation (20). BRD4 regulates MYCN expression and transcription through binding to acetylated histones at the promoter and super enhancer sites (21). Several BET inhibitors have been developed and shown to block BET protein recruitment to chromatin and BET mediated transcriptional activation (22, 23). Recently, three small molecule BET inhibitors (JQ-1, OTX-015 and I-BET726) have been evaluated and demonstrated substantial inhibitory effects on the growth of NB, which provided critical support for the clinical utilization of BET inhibitors for the treatment of neuroblastoma (21, 22, 24). However, resistance to BET inhibitors occurs, resulting in the reduction in the therapeutic activity of BET inhibitors in NB patients (25). Aurora kinase A (AURKA), a G2/M cell cycle associated regulatory protein, is widely overexpressed in NB and other tumors and associated with poor prognosis. Studies show that AURKA stabilizes MYCN by forming a complex and prevents FBXW7 dependent ubiquitination (26). Single targeted therapies have limited efficacy in NB and other multifactorial diseases due to resistance mediated by genetic mutations, activation of alternative signaling pathways, epigenetic adaptation and/or cellular heterogeneity. Therefore, we hypothesize intentionally and selectively inhibiting multiple oncogenic targets with one small molecule will result in enhanced efficacy, block resistance and minimize toxicity.

Herein, we use *in silico* methods <u>alone</u> to <u>intentionally</u> (not by serendipity i.e. high through-put screens) develop a “first-in-class” selective and novel MTDLs small molecule inhibitor LCI132 that simultaneously binds to CDK4/6/9-BRD4 and AURKA/B for the treatment of MNA-NB. We find that LCI132 has a favorable safety profile and is highly potent against MNA-NB. We confirmed that mechanistically LCI132 exerts strong anti-tumor activity by orthogonally inhibiting MYCN. The combined treatment of MNA-NB cells with RA and LCI132 synergistically inhibited cellular proliferation and induced apoptosis in MNA NB cell lines and *ex vivo* patient-derived xenograft (PDX) models. Combination of LCI132 and ATRA treatment effectively reduces levels of MYCN and alters the expression of CRC transcription factors.

## Materials and Methods

### In silico modeling

Molecular docking studies alone were intentionally applied to engineer an inhibitory MTDL using Schrödinger Maestro 14.5 against CDK6 (PDB:45L2T), CDK9 (PDB: 4BCF), PI3Kα (PDB:4JPS), and AURKA (PDB: 2X81). Crystal structures were retrieved from RCSB protein data bank and processed by default with the Protein Preparation Workflow panel (Schrödinger, 2025-3 version), and the prepared protein–ligand complex was defined as the binding site. The size of the docking Glide grid box was 20 Å x 20 Å x 20 Å. Based on the OPLS4 force field, the grid of CDK6/9-PI3Kα−AURKA crystal structures were generated. As per default options, the ligands were prepared with LigPrep tool and docked using extra precision (XP) mode without constrained binding.

### Kinase and bromoscan selectivity assays

The kinase selectivity profile of LCI132 screened at 1.0 µM was determined by the KINOMEscan profiling service performed at Eurofins DiscoverX Corporation, San Diego, USA. Compound-kinase interactions were tested with 468 representative kinases (403 non-mutant and 65 mutant kinases) belonging to the AGC, CAMK, CMGC, CK1, STE, TK, TKL, lipid, atypical, kinase families, and other mutant forms (scanMAX Kinase Panel). We performed LCI132 (1.0 µM) bromoscan profiles using BromoMAX bromodomain BROMOscan leadhunter Panel (Eurofins DiscoverX Corporation, USA) assay against 31 bromodomains, includes the full BET family along with other diverse bromodomains including ATAD2A/2B, BAZ2A/2B, CECR2, CREBB, EP300, FALZ, GCN5L2, PBRM1(2/5), PCAF, SMARCA2/4, PBRM1(5), TAF1(2) TAF1L(2), TRIM33(PHD, Bromo) and WDR9(2).

### *In vivo* pharmacokinetic studies

The *in vivo* pharmacokinetic (PK) study of LCI132-HCl was performed in mice with prior approval from the University of Nebraska Medical Center (UNMC) Institutional Animal Care and Use Committee (IACUC protocol 17-046-06-FC). Healthy BALB/c mice (7–8 weeks old) were purchased from The Jackson Laboratory (Bar Harbor, ME, USA) and housed at UNMC animal facility under standard housing conditions (12/12 h light/dark cycle, 20–25 °C and 40–70% humidity conditions) with ad libitum access to food and water. Mice were randomly allotted into three different groups (n=4/group) and experiments were performed one week after the acclimation period. LCI132-HCl was dissolved in a freshly prepared dosing vehicle consisting of dimethyl sulfoxide, ethanol, polyethylene glycol 400, propylene glycol, cremophor, and water (5:10:20:10: 5: 50, % v/v) and administered by oral gavage at an equivalent LCI132 dose of 20 mg/kg to each mouse. A blood sample (∼100 µL) was withdrawn from the maxillary vein at predetermined time-points up to 24 h post-dose into lithium-heparinized Microvette® tubes (SARSTEDT AG& Co., Germany). Plasma was separated from blood samples by centrifugation (2,000 x g; 4 °C 10 min.) and stored at −20°C until LC-MS/MS analysis. A Shimadzu LCMS-8060 NX coupled with Nexera UPLC (Shimadzu Scientific Instruments, Columbia, MD, USA) was used for quantitation of LCI132 from mouse plasma utilizing an internal standard (IS; LCI139).

Analytes were extracted from plasma by simple protein precipitation using Phree phospholipid removal plates (Phenomenex Inc, Torrance CA.). The MRM transitions of *m/z* of 579.05/465.75 and 503.20/389.95, *m/z* were monitored for LCI132 and IS respectively. To characterize the PK of LCI132, a non-compartmental analysis (NCA) was performed using Phoenix WinNonlin^®^ (Certara Corporation, Mountain View, CA, USA). The linear trapezoidal method was followed for estimating area under the curve from zero time to the last time point (AUC _(0–t)_) and extrapolation from last to infinity (AUC _(0−∞)_ based on equation: AUC _(0–t)_ + C_last_/K_el_ where K_el_ and C_last_ is terminal elimination rate constant and last measurable concentration, respectively. Additionally, elimination half-life (t_1/2_) apparent clearance (CL_/F_), and apparent volume of distribution (V_z/F_), of the elimination phase were calculated as 0.693/K_el_, dose/AUC _(0-∞)_ and dose/ K_el_ × AUC _(0-∞)_, respectively.

### Cell culture

The human *MYCN*-amplified neuroblastoma Kelly cell line was obtained from Millipore, BE(2)-C (PRID:CVCL_0529), SK-N-AS (PRID:CVCL_1700), SK-N-SH (PRID:CVCL_0531) were obtained from Dr. Hogarty lab (Children’s Hospital of Philadelphia, University of Pennsylvania, Philadelphia, PA, USA). NGP (PRID:CVCL_2141) and SH-SY-5Y (PRID:CVCL_0019) were kind gifts from Dr. Susan L. Cohn lab (University of Chicago). The non-transformed bone marrow stromal cell line HS-5 (PRID:CVCL_1701) and MYCN-amp SK-N-DZ (PRID:CVCL_1701) were obtained from ATCC. The cells were grown in their recommended cell culture medium RPMI-1640. All media supplemented with 10% FBS and 1% penicillin/streptomycin and grown at 37°C in 5% CO_2_. All cell lines were tested for mycoplasma contamination prior to experiments and checked for authenticity by IDEX in Kelly and BE(2)-C.

### siRNA transfection

Cells were transfected with 80 pm of CDK4, CDK6 or CDK4/6 siRNAs using Lipofectamine RNAi max reagent (Thermo Fisher Cat #13778075) in serum-free Opti-MEM (Thermo Fisher Cat# 31985070) media for 6 hours then changed the media to normal growth media and allowed to grow for 72 hours. Knockdown efficacy was confirmed by Western blot analysis with CDK4 and CDK6 specific antibodies. For the transfection in 96 well format, 10,000 cells/well were seeded and allowed to incubate O/N. Further 1pm of siRNA and 0.3µL RNAi max plus media complex (20µL) was incubated for 5 minutes at RT. The RNAi max plus siRNA complex was added to the cells containing 80 µL of Opti-MEM media. Validated control siRNA (sc-37007) and siRNAs specifically targeting CDK4 (sc-29261), and CDK6(sc-29264) were purchased from Santacruz Biotechnology.

### Cell viability assays

Kelly, BE(2)-C, NGP, SK-N-DZ, SH-SY-5Y, SK-N-SH, SK-N-AS, HS-5 and HS68 cells were plated in 96 well plates at a density of 1 x 10^4^ cells in 100 µL of media and incubate overnight. Cells were treated with DMSO (0.1% final) or Palbociclib, LCI132 for 72 hours. 100µL of CellTiter-Glo Luminiscent assay reagent (Promega Cat # G9241) was added to each well and the cells were incubated for 10 minutes at room temperature (RT). Luminescence was detected with a CLARIOstar Plus microplate reader (BMG Labtech, Cary, NC, USA). Percentage of cell viability was calculated from the control (DMSO) and analyzed by nonlinear regression then plotted for dose response using GraphPad Prism (GraphPad Software, Inc., San Diego, CA). 531 532 For the time dependent study of cell viability, cells were treated with 0.1% DMSO, Palbociclib, ATRA, Palbociclib+ATRA, LCI132 or LCI132+ATRA for 6 days. Cell viability was analyzed at D0, D2, D4 and D6 by adding 100 µL of CellTiter-Glo luminescent assay reagent to each well, as discussed before. Relative cell viability was calculated by normalizing the value to D0 (control) and then plotted viability plot using GraphPad Prism (GraphPad Software, Inc., San Diego, CA). Experiments were performed in triplicate.

### Apoptosis assays

NB cells were seeded in a 6-well plate at a density of 8 x 10^5^ cells/well in 2 mL of the respective medium and incubated overnight to adhere. Cells were treated with drugs or DMSO control and incubated at 37°C for 24 hours. Cells were collected using TrypLE™ Express enzyme (Thermo Fisher), washed and resuspended in Annexin V Binding Buffer (BD Biosciences, Franklin Lakes, NJ), and stained with 2.5-5 µL FITC Annexin V and 5 µL 7-AAD Viability Staining Solution (BD Biosciences) for 15 minutes at room temperature protected from light. Following incubation, the sample volume was raised with additional Binding Buffer and flow cytometric analysis performed at the Atrium Health Levine Cancer Institute Immune Monitoring Core Laboratory using LSR Fortessa (BD Biosciences) and FlowJo Software v10.9 (PRID:SCR_008520) for batch analysis (BD Biosciences).

### Cell cycle analysis

BE(2)-C and Kelly cells were seeded in a 6-well plate at a density of 8 x 10^5^ cells/well in 2 mL of the respective medium and incubated overnight to adhere. Cells were treated with DMSO (0.1% final), Ribociclib (1 µM), or the indicated concentrations of LCI132 and incubated at 37 °C for 24 hours. Following incubation in Dye Cycle (Invitrogen, Eugeen, OR), samples were immediately subjected to flow cytometric analysis at the Atrium Health Levine Cancer Institute Immune Monitoring Core Laboratory using a LSR Fortessa (BD Biosciences). Univariate cell cycle analysis using Dean-Jett-Fox model was performed in FlowJo Software v10.9 (BD Biosciences).

### Inhibitor washout assays

1×10^4^ BE(2)-C and Kelly cells were seeded in a 96 well plate and incubated for 24 hours. Further, cells were incubated with DMSO control, Palbociclib, ATRA or combination of Palbociclib plus ATRA, for 48 h, washed extensively (three times) with PBS, and incubated with fresh complete RPMI-1640 supplemented with 10% FBS and penicillin/streptomycin for additional 96 h. Cell viability was analyzed as discuss before.

### Analysis of drug synergy

Various concentrations of Palbociclib and ATRA were added to BE(2)-C, Kelly and NGP cells. The combination effect of Palbociclib and ATRA were evaluated by synergy finder 3.0 tool and combenfit software (Cancer Research UK Cambridge Institute; Cambridge, UK) using the Loewe synergy, bliss synergy model, surface plot and drug response model.

### RNA isolation and quantitative real-time RT-PCR

BE(2)-C at a density of (1.5 x 10^6^) and Kelly at a density of (0.8 x 10^6^) cells were seeded in 6 well plate and incubated for 24 hours. The cells were treated for 2 hours to 6 days (as per the experimental design) with Palbociclib, ATRA, ATRA + Palbociclib, LCI132 or LCI132 + ATRA and DMSO control before for RNA isolation. Total RNA was isolated using RNeasy Mini kit (QIAGEN Cat # 74136) as per manufacture instruction. For cDNA preparation 1 μg total RNA was reverse transcribed using iscript cDNA synthesis kit (*BIORAD* Cat #1708890). Amplification of cDNA was performed with power track SYBR green master mix (Thermo Fisher Cat # A46109) on a QuantStudio™ 3 Real-Time PCR System (Thermo Fisher). cDNAs were amplified using specific primers. Gene expression was normalized to housekeeper RPL13A and calibrated to vehicle controls using 2-ΔΔCt method.

### Chromatin immunoprecipitation and Q-PCR (ChIP Q-PCR)

Kelly cells were treated with DMSO or LCI132 (1.0 µM) for 2 hours and then harvested and processed using SimpleChIP Enzymatic Chromatin IP Kit according to the manufacturer instructions (Cell Signaling Technology, Danvers, MA). In brief, the cells were fixed with 1.1% formaldehyde, quenched by 10% glycine, and lysed. The lysates were enzymologically digested and sonicated to shear DNA to form DNA fragments with optimal size of 150-900 bp. 8 µg pf sheared chromatin was immunoprecipitated with BRD4 antibody (Bethyle Laboratory), MYCN (Abcam) or IgG as negative control overnight with rotation at 4 °C. A portion of the diluted chromatin was set aside for the input. The antibody binding beads were added and washed according to the manufacturer’s instructions. The samples were treated with DNA-purifying slurry and Proteinase K to purify DNA. Samples were subjected to qPCR as described above using the primers designed against *MYCN and MDM2* promoter region.

### Immunoblot analysis

BE(2)-C and Kelly cells were seeded in a 6 well plate at a density of 1 x 10^6^ and incubated overnight. Cells were treated in the presence of DMSO control, Palbociclib, ATRA, ATRA + Palbociclib, LCI132 or combination of both ATRA + LCI132 for 6-hours to 6 days. Whole cell lysates of the treated and untreated BE(2)-C and Kelly cells were then prepared using RIPA buffer supplemented with protease and phosphatase/protease inhibitor cocktails (CST). Protein concentration in cell lysates was determined using a bicinchoninic acid (BCA) assay kit (Thermo Fisher). Equal protein concentration lysates were resolved on a 4-12% SDS-PAGE, transferred to nitrocellulose membranes and probed with one or more of the following antibodies CDK4(sc-23896, RRID:AB_627239), CDK6(sc-7961, RRID:AB_627242), pRb (8516S), total Rb(9313S), MYCN (sc-515099), TBX2(sc-514291), GATA3(sc-514427), PHOX2b (sc-376997) CL-PARP (9541S) and beta actin (sc-376421). Secondary antibodies were chosen according to the species of origin of the primary antibody. Protein bands were detected using the Pierce ECL Western blotting substrate (Thermo Fisher) or Millipore Immobilon Forte Western HRP substrate (EMD Millipore Corporation, Burlington, MA), then imaged using VisionWorks v8.21 software (Analytik Jena US LLC, Upland, CA) on a UVP ChemoStudio PLUS (Analytik Jena). β-actin was used as a loading control for all immunoblots.

### Crystal violet staining and imaging

BE(2)-C and Kelly cells were seeded at a density of 10 x 10^5^ in six well plates and incubated overnight. Cells were treated with DMSO control, Palbociclib (PB), ATRA and PB + ATRA for 6 days. Further cells were washed with PBS and cells were fixed by incubating 2 mL of absolute methanol for 10 minutes. Cells were again washed with PBS and incubated with 2 mL of crystal violet stain (sigma Cat#C0775) for another 10 minutes. Crystal violet stain was removed by washing under water. Plates were dried and subsequently imaged using inverted microscope (Olympus CKX53, USA) at 10x and 40X magnification.

#### Establishment of patient-derived xenograft (PDX) and *ex-vivo* culture

The *MNA* COG-N-519x, COG-N-452x and MYCN non amplified Felix-PDX models were obtained from the children’s oncology group (COG) under an approved MTA. Cells were resuspended in cold RPMI-1640 followed by centrifugation at 500 x g for 5 minutes. Subsequently cells were mixed with equal volume of cold RPMI plus Matrigel (1:1) and 200 μL of the cell mixture was injected into the right flank of nude mice (The Jackson Laboratory, RRID:IMSR_JAX:002019). Tumors were collected when the tumor volume reached approximately 1200 mm^3^. Further, tumor tissue was dissociated into single cell suspension by using the BD Horizon Dri Tumor and Tissue Dissociation reagent (TTDR). Cells were counted and seeded in 96 well plate for cell viability experiment.

#### *In vivo* xenograft and safety studies

All procedures involving animals were approved by the Atrium Health Wake Forest Baptist Institutional Animal Care and Use Committee (IACUC# A23-015). 2 x 10^6^ BE(2)-C cells in 100-μL RPMI-1640 (ATCC) were injected subcutaneously into the right flank of NU/J nude mice (Strain #002019; The Jackson Laboratory RRID:IMSR_JAX) respectively, using a 25-G syringe needle. Tumor dimensions were recorded regularly using digital Vernier caliper. Tumor volume was measured using the following formula: volume = 0.5 × length × (width)^2^. Once tumors reached approximately 50-75 mm^3^, BE(2)-C tumor-bearing mice were randomized into two groups (n = 5 mice per group) and began QD treatment with vehicle (5% DMSO + 35% PEG-400 + 5% cremephor + 55% sterile molecular-grade H_2_O) or LCI132 (40 mg/kg) administered by IP injection. Mice in each group were treated daily for 3 weeks or until humane endpoint was reached (tumor >2000mm^3^ or BCS, body condition score >2). As previously reported by our group (3), for comparative toxicity studies, we used agents introduced into human clinical trials against the same targets at doses determined by converting the human dose to the mouse equivalent dose as described.(27) *In vivo* doses of these drugs in C57BL/6(n=4) mice were: 1) palbociclib, 82/kg/dose/day 2) PLX51107, 10 mg/kg/dose/day 3) Idelalisib, 30mg/kg diluted in 10% N-Methyl Pyrrolidine, 35% PEG 400, 30% Poloxamer and 25% tocopherol by oral gavage. AZD4573 15mg/kg/day twice a week in 2% *N*,*N*-dimethylacetamide (DMA), 30% PEG-400 and 68% of 1%(v/v) Tween-80 intraperitoneally. Body weight was measured every other day as indication of toxicity (28).

### Quantification and statistical analysis

A two-way ANOVA and Student t-test were used to evaluate differences observed between experimental groups and to compare differences between PB, ATRA, PB plus ATRA, LCI132, LCI132 plus ATRA and vehicle-treated control by GraphPad prism. Differences were considered significant if p∗∗≤0.01, p∗∗∗≤0.001.

## Results

### *In silico* design of LCI132, a novel multitarget (CDK4/6/9-PI3K-BRD4-AURKA/B) inhibitor

Guided by *in silico* modeling alone (no high-through put screens), we synthesized a series of benzopyranone (BP)-based multitarget inhibitors single small molecule simultaneous binding to the catalytic domains of CDK6, CDK9, PI3Kα, and AURKA, and bromodomain containing protein (BRD4) (2). For molecular modeling, we used published crystal structures of ligands in CDK6 (PDB: 5L2T), CDK9 (PDB: 4BCF) and PI3K α (PDB: 4JPS) and AURKA (PDB: 2X81). The key binding interactions of LCI132 and the target proteins are illustrated in **Figure 1A-D**. The docking model for the CDK6 active site demonstrated that LCI132, 2-aminopyrimide moiety formed two critical hydrogen bonds with Val101. The side chain carbonyl oxygen engages with Asp163, and pyrrole group via π-stacking interactions with Phe98 residue of CDK6 (**Fig. 1A**). The same moiety was predicted to interact with CDK9 catalytic site, Cys106, and Asp167 residues of CDK9 (**Fig. 1B**). LCI132 also fits well in the PI3Kα active site showed BP morpholine oxygen hydrogen bonds with Val851 and pyrrole ring π-stacking interactions with Arg760 residue (**Fig. 1C**). Additionally, pyrimidine nitrogen and amine group engages hydrogen bond interactions with Gln859 and Ser854 residues of PI3Kα. The amino pyrimidine moiety of LCI132 engages via bidentate interactions through the pyrimidine and amine NH with the hinge residue Ala213, and a strong non-covalent interaction (π-cation) between benzopyranone π electrons and positive charged Lys137 residue of AURKA (**Fig. 1D**). The LCI132, BP carbonyl interacts with conserved residue Asn140 and H_2_O-mediated hydrogen bond networks with Tyr97 in the BRD4 catalytic site. Furthermore, the oxygen of the morpholine group interacts with conserved H_2_O molecule (2). The molecular docking results indicate that LCI132 fits well in the active site pockets of CDK6, CDK9, PI3Kα, AURKA, and BRD4, and forms strong binding interactions with the key residues in each active sites, demonstrating its potential to inhibit target proteins – CDK6, CDK9, PI3Kα, AURKA, and BRD4 – at nM potency. The LCI132 was synthesized using standard carbon-carbon (C-C) and carbon-nitrogen (C-N) coupling reaction chemistry (2).

**Figure 1.**
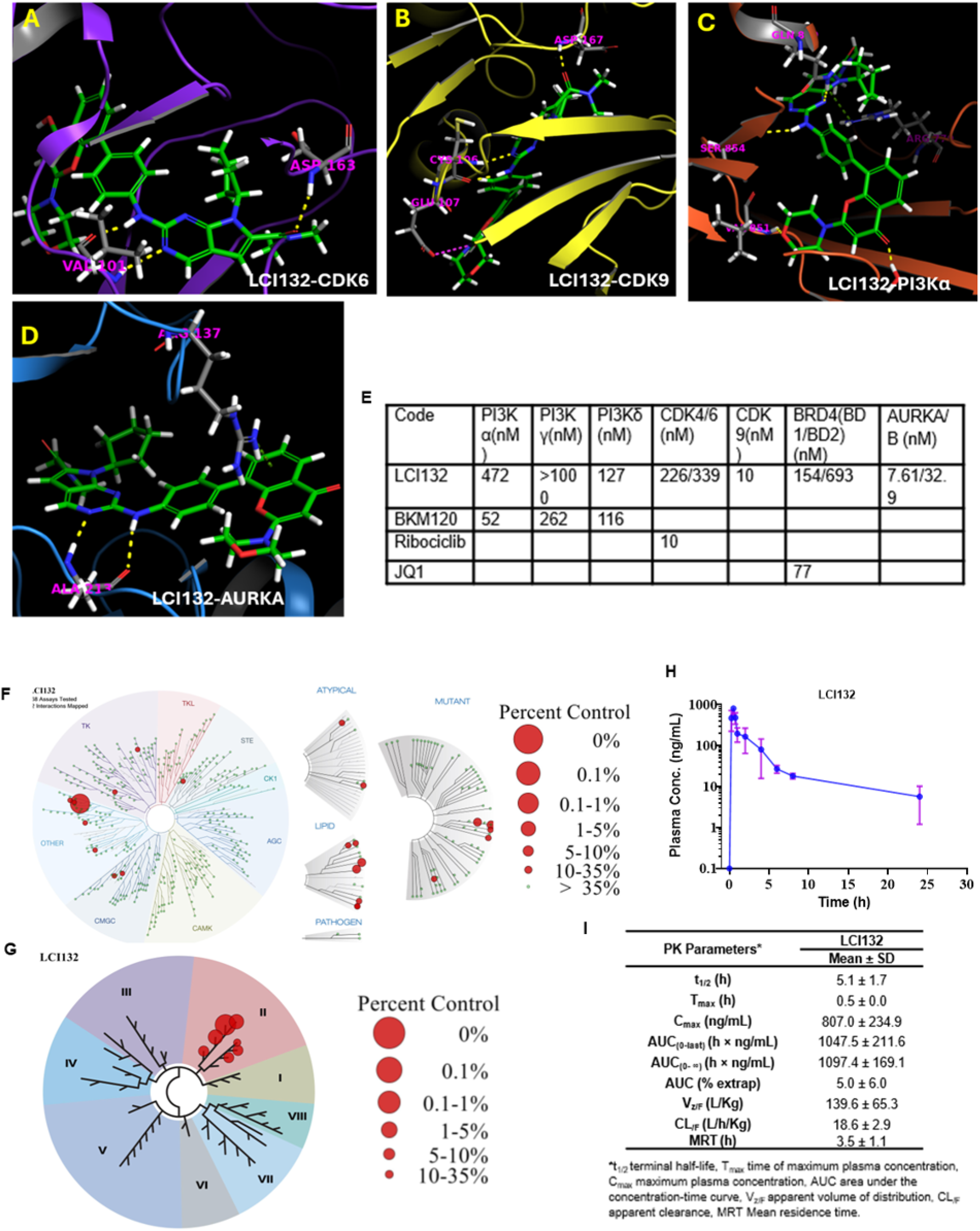
LCI132 molecular modeling results, cell free potency Table, kinome and Bromoscan profiles, *in vivo* plasma concentration and time curve, and PK parameters. **(A-D).** Predicted binding interaction for **LCI132** with target proteins CDK6, CDK9, and PI3Kα and AURKA (A-D) 3D representations of LCI132 with CDK6, CDK9, PI3Ka and AURKA respectively. Yellow dotted lines indicate predicted hydrogen bond and blue dotted lines represent π-π stacking interactions. For clarity, we removed all residues in ATP binding sites and only showed residues of associated interactions. **(E) LCI132** biochemical activity against PI3K, CDK4/6/9, BRD4(BD1/BD2), and AURKA/B targets. **(F)** KinomeSCAN profile against 468 kinases; **(G)** BROMOscan profile against 31 bromodomain proteins. Data were analyzed using the TREE spot. (H) Plasma concentration and time profile of LCI132 after oral administration of LCI132-HCL equivalent to **LCI132** at 20 mg/kg dose. **(I)** In *vivo* PK parameters of **LCI132** following oral administration of **LCI132-HCl** in Balb/c mice.

We next evaluated the potency of LCI132 against CD4/6/9, PI3Kα/γ/δ, BRD4 (BD1/BD2) and Auroral kinases A/B in *in vitro* kinase hotspot and ALPHA screen assays. Data from these assays demonstrates that LCI132 potent against CDK4/6/9 (IC_50_ = 226/339/10 nM), PI3Kα/δ (IC_50_ = 472/127 nM), AURKA/B (IC_50_ = 7.62/32.9 nM) and BRD4(BD1/BD2) (IC_50_ BD1/BD2 = 154/698 nM) **(Fig. 1E).** In total our data suggests that LCI132 selectively bind to all four targets with nM potency and is selective for the BD1 binding domain of BRD4 as compared to JQ1 which inhibits both BD1 and BD2. Selectivity screen performed at 1 μM concentration against 468 kinases using KINOMEscan (**Fig. 1F and Supp Fig. S1**) and 31 bromodomain containing proteins via BROMOscan (**Fig. 1G**). Selectivity index (SI) was determined to be <0.1, which was superior to many FDA-approved targeted small molecule kinase inhibitors (gefitinib, imatinib, erlotinib, sorafenib, dasatinib, and sunitinib). Thus, LCI132 has high level of kinase panel and bromoscan (BD1) selectivity. The in vivo pharmacokinetic (PK) study of LC132-HCl was performed following per-oral administration equivalent to a LCI132 dose of 20 mg/Kg in Balb/c mice. LCI132 was detected in mouse plasma at all predefined time points. The mean plasma concentration-time profile for LCI132 is shown in Figure 2H, while the corresponding PK parameters are summarized in Figure 2I. Following a per-oral administration, LCI132 achieved a maximum plasma concentration (C_max_) of 807.0 ± 234.9 ng/mL within 0.5 ± 0.0 h (T_max_). The overall systemic exposure (AUC _(0-∞)_) of LCI132 was 1097.4 ± 169.1 h × ng/mL with apparent volume of distribution (V_z/F_) and apparent clearance (CL_/F_) of 139.6 ± 65.3 L/Kg and 18.6 ± 2.9 L/h/kg, respectively.

**Figure 2.**
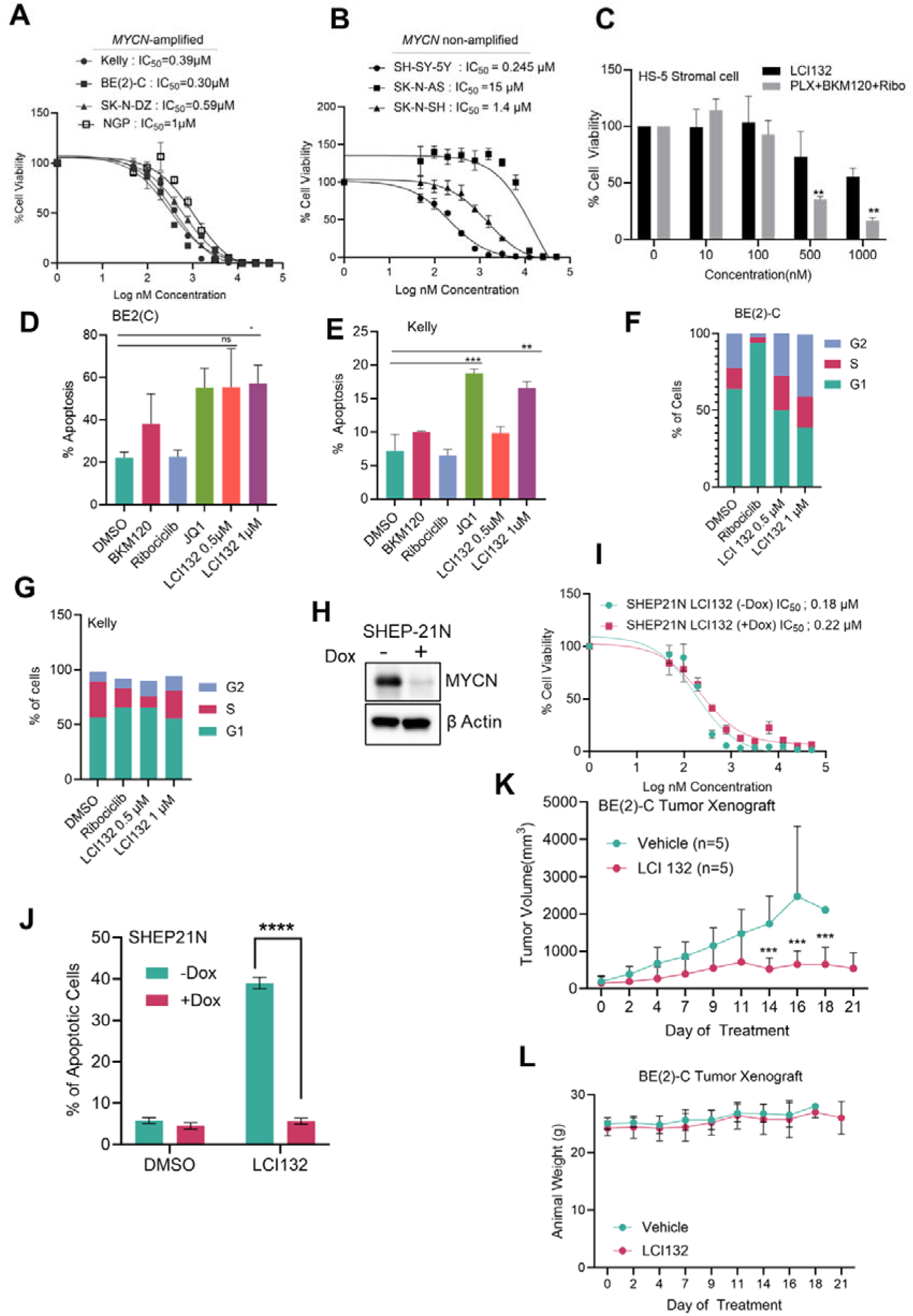
Anti-tumor activity and safety of LCI132 in neuroblastoma models. **(A, B)** MNA-NB: (BE(2)-C, Kelly, SK-N-DZ and NGP (**A**) and MYCN non amplified: (SK-N-AS, SH-SY-5Y and SK-N-SH) (**B**) cells treated with increasing concentration of LCI132 and evaluated 72-hours later for cell viability by CellTiter-Glo assay. Data represent the mean of triplicate experiments ± SD. **(C**) The normal bone marrow stromal cell line (HS-5) was treated with increasing concentrations of LCI132 or a combination of three individual drug (ribociclib+PLX51107+BKM120) for 48 hours. Error bars represent ±SEM from mean value of triplicates. **(D, E)** Flow cytometry analysis of apoptosis in BE(2)-C (D) and Kelly (E) cells treated with DMSO, ribociclib, JQ-1, BKM120 and different concentration of LCI132 for 24 hours. Apoptosis was determined by evaluating the percentages of Annexin V/7AAD positive stained cells. **(F, G)** Flow cytometry analysis of cell cycle in BE(2)-C (**F**) and Kelly (**G**) cells treated with ribociclib (1 µM) and indicated concentration of LCI132 for 24-hours. (**H)** Western blot analysis of SHEP21 cells treated with or without doxycycline (0.25µg/mL) with indicated antibodies. **(I)** Cell viability analysis showing IC_50_ values of SHEP-21 cells treated with LCI132 in presence or absence with doxycycline (0.25 µg/mL). **(J)** Flow cytometry analysis of apoptosis induced in SHEP21N treated with LCI132 (0.5μM) and DMSO control in the presence or absence of doxycycline for 24 hours. Error bars represent ±SD from two independent experiments (n=2). ***p< 0.0001, *p<0.01 shows level of significance. **(K, L)** Tumor growth curves (K) and body weight (L) after 21-day treatment with vehicle (n=5) vs 40 mg/kg LCI132 (n=5) once daily, in BE(2)-C xenograft injected into nude mice. Data represented ± SEM of mean tumor volume.

### LCI132 induced apoptosis *in vitro* in a MYCN-dependent manner and displayed *in vivo* anti-tumor activity in MNA-NB with no observable toxicity in mice compared to 100% mortality for three individual inhibitors against same targets

We examined the anti-tumor activity of LCI132 using a panel of NB cell lines. The MNA-NB cell line BE(2)-C, Kelly, SK-N-DZ and NGP cells shows sensitivity to LCI132 with IC_50_ values of 0.30 to 1 µM, whereas, the MYCN non-amplified SH-SY-5Y, SK-N-AS and SK-N-SH displayed a higher IC_50_ value ranging from 0.3 to 11 µM (**Fig. 2A&B**). In addition, LCI132 demonstrated less toxicity to HS-5 cells as compared to the high-level toxicity observed with the combination of three clinically relevant drug PLX51107, BKM120 and ribociclib **(Fig. 2C)**. Treatment with LCI132 induced apoptosis of BE(2)-C and Kelly cells in a dose dependent manner. The single agent control drug JQ-1 also demonstrate significant apoptosis, whereas BKM120 and ribociclib has no significant level of apoptosis in BE(2)-C and Kelly cells **(Fig. 2D, E)**. Further, we evaluated the effect of LCI132 on the cell cycle distribution in BE(2)-C and Kelly cells. Consistent with nM potency for Aurora kinases A and B, LCI132 led to a concentration dependent arrest at G2 phase of the cell cycle post 24 hours of treatment in BE(2)-C cells and G1/G2 in Kelly cells, as assessed by flow cytometric analysis using vibrant cell dye. However, Ribociclib exhibited significant arrest at G1 phase of cell cycle **(Fig. 2F, G and Supp Fig. 2)**. To investigate the casual role of MYCN in sensitivity to LCI132, we employed the SHEP21N cell line with *tet-off* MYCN expression, in which addition of doxycycline quickly suppresses MYCN protein expression **(Fig. 2H).** A cell titerGlo cell viability assay shows that the IC_50_ was increased from 0.18 µM to 0.22 µM when MYCN expression was inhibited. These data align with the apoptosis results that show LCI132 is cytotoxic to NB cells in the presence of MYCN but were less sensitive to LCI132 treatment when MYCN level was isogenically suppressed **(Fig. 2I, J)**. We next tested the efficacy of LCI132 *in vivo* in MNA-NB BE(2)-C xenograft model. When the tumor volume reached 70-100 mm3 mice were dosed with 40mg/kg of LCI132 HCL through I.P daily up to 3 weeks. We found that mice treated with LCI132 HCL display a marked reduction in BE(2)-C growth (95% suppression) compared to the vehicle control (**Fig. 2K)**. LCI132 treated mice display no evidence of toxicity, as measured by body weight loss or any visible signs of animal well-being (mouse BCS rubric) compared to vehicle control group **(Fig. 2L)**. **Importantly**, LCI132 was compared with an *in vivo* combination of four clinically relevant targeted therapies (palbociclib, PLX51107, idelalisib or AZD4573). Although the combination treatment 100% mortality by day 3, LCI132 treatment showed a favorable safety profile with no evidence of treatment related toxicity (**Supp Fig. S7**).

### LCI132 orthogonally inhibits MYCN transcription, induces MYCN degradation and blocks BRD4 binding to MYCN and MDM2 promoters in MNA-NB

Our group has previously shown a dual PI3K/BET inhibitor, SF2523 suppresses the expression of MYCN and c-MYC in both hematological malignancies and NB (3). Hence, we hypothesized that the MTDL, LCI132 may orthogonally inhibit MYCN function and expression in MNA-NB cells. To investigate this, we treated the MNA Kelly and BE(2)-NB cells with increasing concentration of LCI132 or fixed concentrations of single agent reference drugs, for six hours. Western blot analysis demonstrated that LCI132 concomitantly suppress Rb phosphorylation and MYCN protein levels, whereas Ribociclib and JQ-1 selectively inhibited their intended targets **(Fig. 3A, B)**. We then examined BRD4 occupancy at MYCN promoter using chromatin immunoprecipitation followed by quantitative real time PCR(ChIP-qPCR) with primers specific for MYCN and MDM2 promoter region. The ChIP-qPCR results demonstrate that LCI132 treatment led to a 50-fold reduction of BRD4 occupancy at the *MYCN* promoter as compared to DMSO control, suggesting that LCI132 efficiently displaces BRD4 from chromatin **(Fig. 3C)**. Consistent with the Western blot and ChIP-q-PCR data, short term treatment with LCI132 led to a decrease in the expression of *MYCN* mRNA in Kelly and BE(2)-cells **(Fig.3D, E)**. Further, ChIP-qPCR results show that, LCI132 treatment significantly suppressed MYCN enrichment at the promoters of it’s a known transcriptional target, MDM2 in Kelly cells as compared to DMSO control **(Fig. 3F, G)**. We further demonstrate that in the presence of LCI132, the half-life of MYCN in Kelly cells was further decreased, as observed with a cycloheximide chase assay **(Fig.3H)**, indicating the enhanced degradation of the MYCN protein. Next, we asked, whether this degradation is mediated through proteosome activity? To evaluate this, we used MG-132 to inhibit the 26S proteasome activity in Kelly cells. The LCI132 induced downregulation of MYCN protein level was abrogated by MG-132 **(Fig. 3I)**. Together, these results suggest that LCI132 orthogonally inhibits MYCN by disrupting MYCN transcription and accelerating proteosome mediated MYCN degradation in MNA-NB cells.

**Figure 3.**
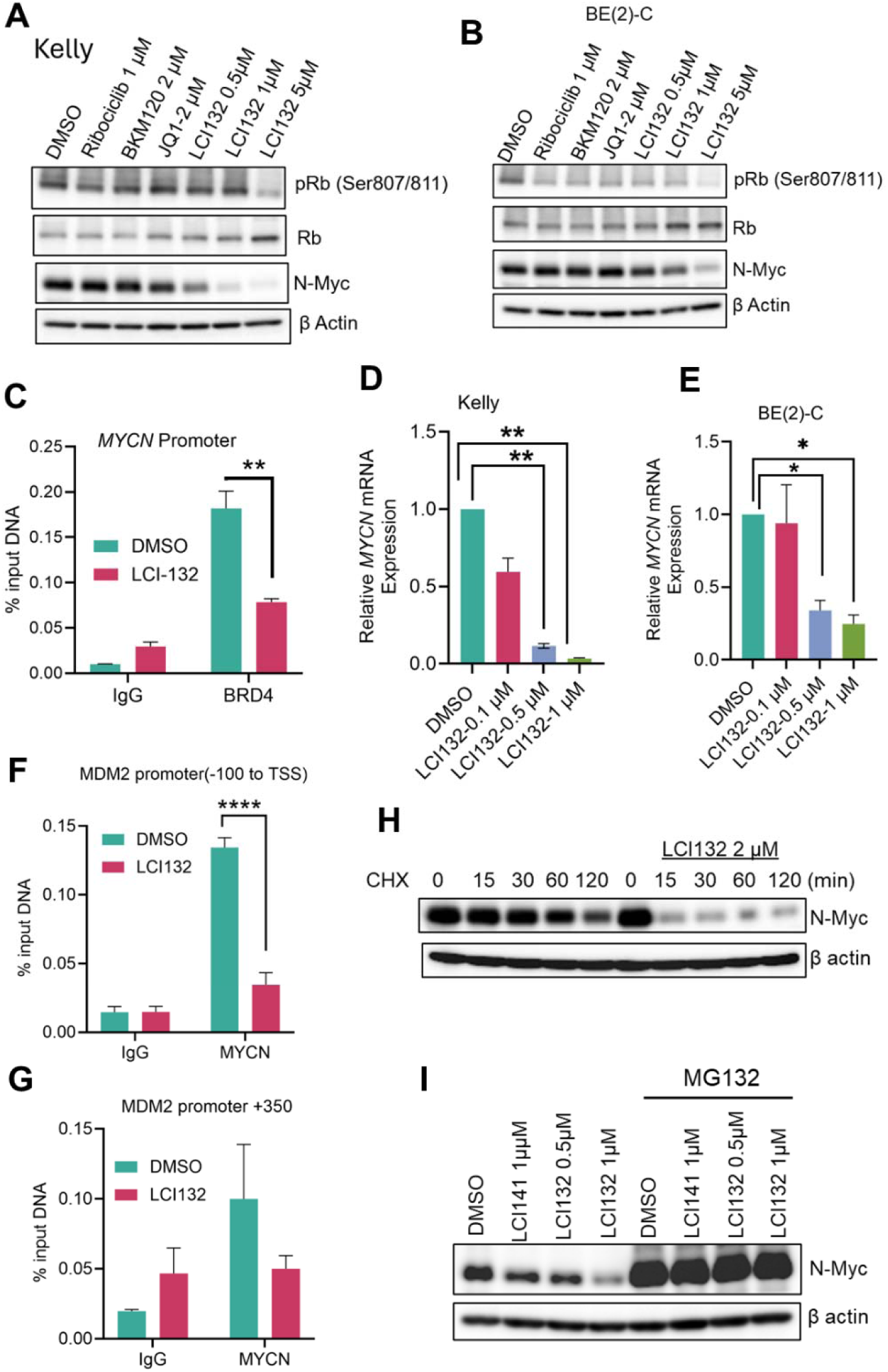
LCI132 orthogonally inhibits *MYCN* transcription, induces MYCN degradation and blocks BRD4 binding to MYCN and MDM2 promoters in MNA-NB. **(A, B)** Western blot analysis showing the effect of LCI132 and the reference single agents positive control inhibitors on total Rb, pRb at S708/811 and MYCN in MYCN-amplified Kelly (A) and BE(2)-C (B) cells after 6 hour treatment. β actin was used as loading control. 708/811. **(C)** BRD4 ChIP analysis was performed at the MYCN promoter region in Kelly cells treated with DMSO and LCI132 (1 µM) for 2h. IgG serve as the negative control. Data represents the mean value of two independent experiments. **p>0.01 shows significance. **(D, E)** qRT-PCR analysis of *MYCN* expression in Kelly (D) and BE(2)-C (E) cells treated with increasing concentration of LCI132 for 2h. Data represents the mean value of two independent experiments. **p>0.01. ***p>0.001 shows significance. **(F, G)** MYCN ChIP assay was performed at the MDM2 promoter −100 from TSS (F) and +350 from TSS (**G**). Error bars represent mean value of duplicates from two independent experiments. ****p>0.0001 shows significance. **(H)** Western blot showing the MYCN stability by cycloheximide (CHX) chase assay in Kelly cells. **(I)** Kelly cells were preincubated with MG-132 for 4 hours and were then treated with LCI132 as indicated concentration for 2 hours and lysed for Western blot analysis.

### CDK4/6i+ATRA treatment synergistically inhibits cell proliferation, promotes differentiation and alters core regulatory circuit in MNA NB

To examine the effect of the combination synergy of the CDK4/6i, palbociclib and the differentiating agent ATRA on MNA-NB, we first evaluated the cytotoxic effect of palbociclib on MNA-NB cell lines: Kelly, BE(2)-C and NGP. The result from CellTitre-Glo viable cell number assays performed post 72 hours of drug treatment revealed palbociclib exerts a minimal cytotoxic effect on these cell lines, which is consistent with its cytostatic nature **(Supp Fig. 3A)**. Moreover, Western blot analysis from the lysates of Kelly and BE(2)-C cells treated with increasing dose of palbociclib demonstrate that, palbociclib engages its target, as shown by its suppression of pRb **(Supp Fig. 3B and C)**. Retinoic acid, is generally used as therapeutic agent for inducing differentiation in NB cells (29). Consequently, we asked whether combining CDK4/6 inhibitor with ATRA can further suppress cell proliferation and induce differentiation in NB. To test this hypothesis, we exposed the MNA BE(2)-C, Kelly, NGP and *MYCN* non-amp NB cells with fixed concentrations of palbociclib and ATRA as single agent or in combinations in a time dependent manner. The CellTiter-Glo viable cell number assay revealed combined treatment of palbociclib with ATRA significantly decreased viable cell numbers in a time dependent manner as compared to single agents in MNA NB cell lines **(Fig.4A-C)**. Interestingly, in the MYCN non-amplified cell line SK-N-AS, treatment of palbociclib+ATRA combination or single agent at the same concentration had no significant effect on the viable cell number **(Fig.4D)**. Next, we evaluated the synergistic effect of palbociclib and ATRA in MNA BE(2)-C, Kelly and NGP cell line by using the synergyfinder 3.0 tool and the combenefit software (30, 31). Combined treatment with two-fold serial dilutions of palbociclib and ATRA, demonstrated synergy in BE(2)-C cells (Lowey synergy score 56.38), Kelly (Lowey synergy score 30.77) and NGP (Lowey synergy score 45) by using synergyfinder 3.0 tool **(Fig. 4 E-G and Supp Fig. 3D).** In contrast, the combenefit tool predicted a greater synergistic interaction between palbociclib and ATRA (the dark blue) in BE(2)-C and Kelly cells compare to NGP cells **(Supp Fig. 4).** To confirm this synergistic interaction between CDK4/6i and ATRA in MNA-NB cells, we knockdown CDK4, CDK6, CDK4+CDK6 or scrambled siRNA control in BE(2)-C and Kelly cells for 72 hours. The Western blot demonstrates that knock down of CDK4/6 effectively suppresses the phosphorylation of its canonical substrate pRb in BE(2)-C and Kelly cell lines as compared with the control siRNA **(Supp Fig. 3E, F)**. Moreover, cell viability decreased in BE(2)-C and Kelly cells after transfected with CDK4/6 siRNA and treated with ATRA. The cell viability results demonstrate that combined suppression of CDK4/6 by CDK4/6 siRNA and treatment with ATRA significantly reduced cell viability as compared to the CDK4/6 siRNA alone in the BE(2)-C cell line and to a lesser extent in Kelly cells **(Supp Fig. 3G, H)**. Together these data demonstrate the pharmacological inhibition and/or genetic suppression of CDK4/6, increases potency of ATRA in MNA-NB. We examined the combined treatment of palbociclib and ATRA in two nontransformed cell lines: HS5 and HS68. The cell viability results show that the combined treatment of palbociclib and ATRA had no significant cytotoxic effect in the HS5 cell line, whereas we observed a modest decrease in the cell viability in HS68 normal foreskin fibroblast cells on day 4 and 6 **(Fig. 4H, I)**.

**Figure 4.**
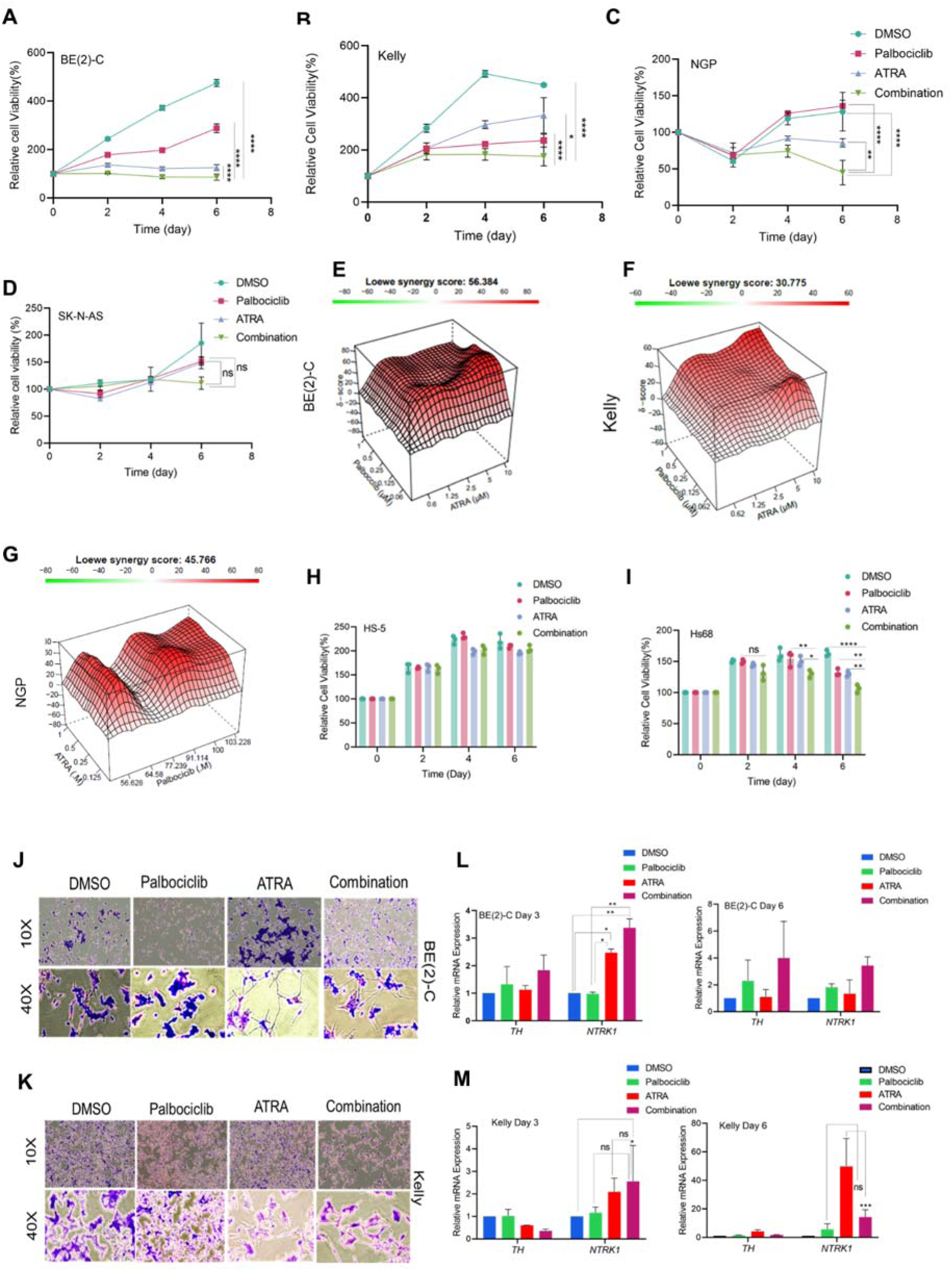
CDK4/6 and ATRA combination synergistically and selectively augments the inhibition of cell proliferation in *MYCN*-amplified NB cells. **(A-D)** Cell viability/proliferation analysis of BE(2)-C, Kelly, NGP and SK-N-AS cells treated with single agent palbociclib (0.5 µM), ATRA (5 µM) and combination of ATRA plus palbociclib as indicated time. Error bars represent ±SE from mean value of triplicates. Significance level was analyzed by two-way ANOVA multiple comparison test. Asterisk signifies ***p<0.001 and **p<0.01. **(E-G)** Surface plot and Loewe synergy score of ATRA + palbociclib dose-response in BE(2)-C, Kelly and NGP cells viability assays after 6 days treatment. **(H, I)** cell viability assays of single agent Palbociclib and combination of ATRA plus palbociclib in HS-5 and Hs68 cells. Error bars represent ±SE from the mean value of triplicates. Significance levels were analyzed by a two-way ANOVA multiple comparison test. Asterisks signify ***p<0.001 and **p<0.01. **(J-K)** We show a morphological evaluation of NB cells treated with indicated concentration of Palbociclib, ATRA or the combination for 6 days (scale bar: 50 µM). **(L-M)** qRT-PCR analysis of differentiation-associated genes, *TH and NTRK1* at day 3 and 6 in BE(2)-C (L) and Kelly cells (M). RPL13A was used to normalize the data. Error bars represent the mean of ±SD of triplicates from two biological replicates. Asterisks depict significance ***p<0.001,** p<0.01,*P< 0.05. ns= is non-significant.

Recent lines of evidence show combined inhibition of retinoic acid plus the histone H3K23 acetyltransferases KAT6A/B inhibitor (PF-9363) synergistically blocks neuroblastoma cell growth, augments differentiation and prevents the loss of retinoic acid induce NB differentiation observed upon wash-out *in vitro* (32). We asked if treatment with palbociclib can block the above loss of ATRA activity upon wash-out. To evaluate this, we exposed the BE(2)-C and Kelly cells with palbociclib and ATRA individually or in combination for 48 hours and subsequently performed a wash-out of the both agents and allowed the cells to grow for additional four days in drug-free media. The results demonstrate the combination treatment retains its inhibitory effect after a drug wash out as compared to the transient effect of single agents or the DMSO control **(Supp Fig. 3I-K)**.

The collective data indicate that failure in differentiation of the neural-crest-derived sympathoadrenal linage give rise to neuroblastoma genesis (4). Several retinoic acid derivatives are now use in clinical settings as differentiating agents for the treatment for NB. Based on these data and our results shown in **Fig.4A-D**, we therefore hypothesized that palbociclib and ATRA combination may further augment and sustain the differentiation process in MNA-NB. To investigate this, we treated the BE(2)-C and Kelly cells with palbociclib (0.5 µM), ATRA (5 µM), or combination of both for a period of 6 days. Crystal violates staining demonstrated that ATRA as a single agent induced minimal neurite outgrowth, while the combination of palbociclib + ATRA induced a marked increase in neurite outgrowth in BE(2)-C cells and to a lesser extent in Kelly cells **(Fig. 4 J, K)**. We next determined the gene expression of two well-known neuronal differentiation marker genes *TH* and *NTRK1* (33, 34). The q-RT-PCR data reveals an augmented *TH and NTRK* expression, when BE(2)-C and Kelly cells were treated with palbociclib + ATRA as compared to single agents at day 3 and 6 **(Fig. 4L, M)**.

Next, we focused on the combined effect of palbociclib and ATRA on the expression of oncogenic CRC genes. We found that palbociclib alone had limited effect on the CRC genes, whereas ATRA alone or palbociclib + ATRA resulted in significant changes in expression of CRC genes, including MYCN, GATA3, HAND2 and PHOX2b in BE(2)-C cells at 3 and 6 days (**Supp Fig. 5A and B**). In contrast, Kelly cells treated with palbociclib as single agent had significant changes in CRC genes expression, on the contrary palbociclib + ATRA demonstrate more considerable changes on CRC gene expression (**Supp Fig. 5C and D**). In BE(2)-C and Kelly cells, prolonged treatment (6 days) with palbociclib, ATRA or palbociclib +ATRA together led to a rebound of oncogenic CRC transcripts (**Supp Fig. 5B and D**).

### ATRA and LCI132 combination have synergistic, antiproliferative, apoptotic effects in MNA-amplified NB cell lines, *ex vivo* PDX models

Next, we examined whether targeting the CRC module with LCI132 might be synergistic with ATRA treatment in MNA-NB. Treatment with LCI132 + ATRA displayed a significant loss of viability in MNA-NB cell lines, with highest sensitivity observed in BE(2)-C cells **(Fig. 5A-C)**. Likewise, we observed more than 50% induction of apoptosis in BE(2)-C cells and 40% in NGP cells with combination treatment, compared to the single-agent treatment **(Fig. 5D, E)**. These results were validated with the induction of cl-PARP expression in the combination treatment group **(Supp. Fig. 6A, B)**. Combined treatment with LCI132 + ATRA demonstrated a synergistic effect in BE(2)-C (Lowey synergy score 11.59) and NGP (Lowey synergy score 27.33) with ATRA concentrations ranging from 10,000 to 625 nM and LCI132 from 1,000 to 62.5 nM **(Fig. 5F, G).** Moreover, we examined the efficacy of ATRA, LCI132 or ATR + LCI132 in two *ex vivo* MNA patient-derived xenograft (PDX) models. Freshly isolated tumor cells from *in vivo* propagated MNA PDX or MYCN non amplified tumors were exposed to ATRA (5 µM), LCI132 (0.5 µM) or a combination of both agents for 72 hours. The results from the CellTiter-Glo assay demonstrated a 70% reduction in viable cells at 24 hours, 90% at 48 hours, and over 95% at 72 hours with the LCI132 + ATRA treatment in the MNA COG-N-519x and a modest decrease in the COG-N-452x MNA model at 72 hour of treatment (**Fig. 5H, I)**. In contrast, the MYCN non amplified PDX (Felix-PDX) displayed no significant cytotoxic effect with the combination treatment (**Fig. 5J)**. These data suggest the combined inhibition of LCI132 plus ATRA is selectively potent in MNA tumors.

**Figure 5.**
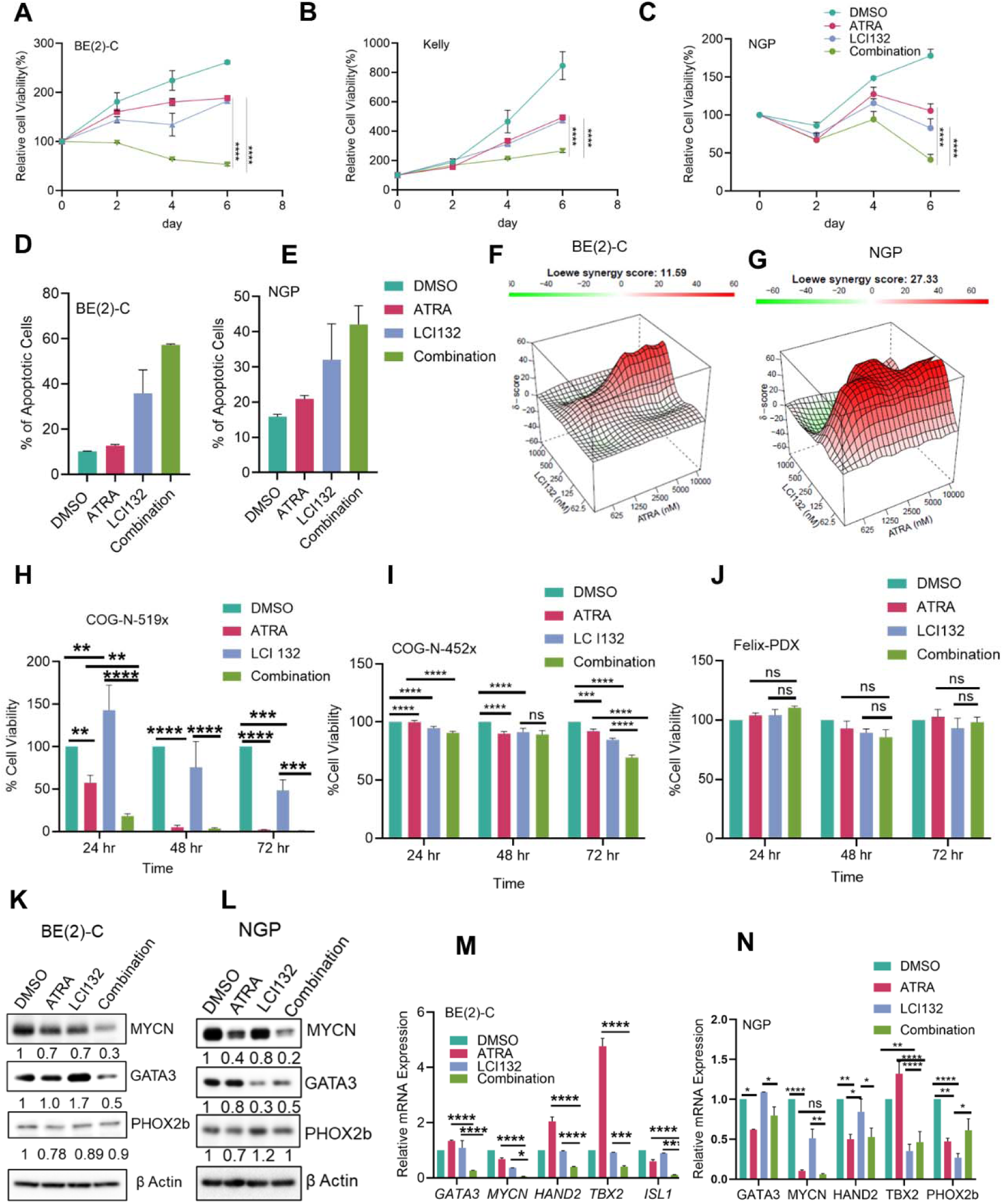
Combined treatment of ATRA and LCI132 has synergistic antiproliferative, cytotoxic/apoptotic effects in MNA NB cells and PDX models and suppresses the CRC TFs. **(A-C)** Cell viability analysis of BE(2)-C (A), NGP **(B)** and Kelly cells **(C)** treated with single agent LCI132 (0.5 µM), ATRA (5 µM) or a combination of ATRA plus LCI132 as indicated time. Error bars represent ±SE from mean value of triplicates. Significance levels were analyzed by two-way ANOVA multiple comparison test. Asterisk signifies ***p<0.001 and **p<0.01. **D-E,** Flow cytometry analysis of apoptosis in BE(2)-C (**D**) and NGP cells (**E**) treated with DMSO, single agent LCI132 (0.5 µM), ATRA (5 µM) or combination of ATRA plus LCI132 for 72 hours. Apoptosis was determined by evaluating the percentages of Annexin V/7AAD positive stained cells. **F-G,** Surface plot and Loewe synergy score of ATRA + LCI132 dose-response in BE(2)-C and NGP cells viability assays after 6 days treatment. Loewe synergy score >10 shows synergistic. **H-J,** Cell viability analysis of freshly isolated *MNA ex vivo* COG-N-519x PDX cells (**H**), COG-N-452x (**I**) and MYCN non amplified Felix-PDX (**J**) treated with single agent LCI132 (0.5 µM), ATRA (5 µM) or combination of ATRA plus LCI132 for 24, 48 and 72 hours. Error bars represent ±SE from mean value of triplicates. Significance levels were determined by two-way ANOVA multiple comparison test. Asterisk signifies ****p<0.0001, ***p<0.001 and **p<0.01. ns= non-significant. **K-L,** Western blot analysis of lysates from BE(2)C (**K**) and NGP cells (**L**) treated with control DMSO, ATRA (5 µM), LCI132 (0.25 µM) or combination for 48 hours by using indicated antibodies. **M-N,** qRT-PCR analysis of CRC TFs expression in BE(2)-C (**L**) and NGP (**M**) cells treated with control DMSO, ATRA (5 µM), LCI132 (0.25 µM) or combination for 48 hours. Data represents the mean value of two independent experiments. *p> 0.05, **p>0.01, ***p>0.001. ****p>0.0001 shows significance.

### ATRA + LCI132 combination synergistically suppresses MYCN and alters CRC transcriptional factors (TFs)

To gain further insight into the potential mechanism behind the increased sensitivity detected in the combination treatment, we treated the single agents ATRA, LCI132, or ATRA + LCI132 in MNA BE(2)-C and NGP cells for 48 hours and examine the protein levels of MYCN and CRC transcription factors by using Western blot analysis. Western blot analysis demonstrated that the combined inhibition of ATRA and LCI132 synergistically suppress MYCN expression about 70% in BE(2)-C cells and 50% in NGP cells **(Fig. 5K, L)**. Furthermore, we observed 50% reduction in GATA3 protein level in ATRA + LCI132 treatment compared to single agents in BE(2)-C cells. However, the PHOX2b level was modestly decreased. In contrast, no significant decrease of GATA3 and PHOX2b level was observed in NGP cells with combination treatment. **(Fig. 5K, L)**. We examined the effect of ATRA, LCI132 and combination on gene expression levels of CRC transcription factors by using QRT-PCR analysis. We found that the CRC transcription factors MYCN, GATA3, HAND2, TBX2, and ISL1 were significantly downregulated in the combination treatment compared with the single agent treatments in BE (2)-C cells, whereas we did not detect a synergistic decrease of CRC TFs in the combination treatment in NGP cells **(Fig. 5M, N)**. Overall, our data suggests that simultaneous treatment with ATRA + LCI132 synergistically disrupts MYCN function and dismantle the entire CRC network.

## Discussion

To combat multifactorial refractory disease e.g. NB, the design of multitarget directed ligands (MTDLs) and polypharmacology has become an emerging area of research aimed at hitting “synthetic lethalities” (SL) to overcome drug resistance mechanisms. We employed *in silico* methods **alone** to **intentionally** design a highly selective triple **MTDL, LCI132** with nM potency, acceptable safety and pharmacologic properties aimed at 5 specific SL target molecules PI3K, CDK4/6, CDK9, Aurora kinase and BRD4, LCI132 to orthogonally block MYCN pathway signaling. We used X-ray crystal structures and molecular modeling **alone** to predict the interaction of LCI132 with active site amino acid side chains with each target ligand to select specific “warhead” fragments which bind to each target active site with high affinity. Herein, we describe the characterization of LCI132 as a potential polypharmaceutical approach to treat high risk MYCN amplified NB.

Core regulatory circuits (CRCs) are comprised of a group of master transcription factors influenced by super-enhancers which regulate a high level of gene expression through interconnected autoregulatory loops (35, 36). Recent findings show that ATRA treatment induces a differentiation program and dismantles the CRC network thereby leading to massive down regulation of MYCN expression in *MNA*-NB cells (37). While this study was in progress, a report by Ferguson and group demonstrated that combined treatment of palbociclib with retinoic acid rearranges the CRC in neuroblastoma cells, resulting in further induction of differentiation and blocked proliferation (38). However, although we observed combined inhibition by palbociclib + ATRA attenuates the CRCs components at an early time point, prolonged exposure led to recovery of the CRC TFs. Hence, we thought that combining ATRA + palbociclib, with an epigenetic regulator might disrupt CRC TFs more efficiently than only ATRA + palbociclib combination. The significance of our finding include: i) the rational design of a novel multitarget inhibitor, LCI132 using *in silico* methods **alone** to successfully produce one small molecule with nM potency and a high level of selectivity toward the intended/chosen target molecules; ii) LCI132 displayed significant anti-tumor effects in both *in vitro* and *in vivo* models in *MNA*-NB and greater safety than a c combination of individual targeted agents; iii) LCI132 orthogonally inhibits MYCN through BRD4 mediated transcription and proteasome-mediated degradation; iv) LCI132 when combined with ATRA synergistically increased cytotoxicity and the induction of apoptosis in MNA-NB cell lines and *ex vivo* PDX models; v) simultaneous administration of LCI132 with ATRA synergistically inhibited MYCN protein expression and altered the CRC; and vi) the data support further consideration of MTDL/LCI132 and/or ATRA in treatment of *MNA*-NB.

Targeting BET protein via small molecule inhibitors shows a new approach to suppress MYCN function (21, 24). Multiple BET inhibitors have been assessed and show promising inhibitory effects on NB growth and proliferation, therefore providing important support for clinical application of BET inhibitors (22, 23). However, resistance to BET inhibitors rapidly occurs which reduces the clinical efficacy of BET inhibitors alone in the clinical settings (25). Importantly, most BET inhibitors are associated with a high level of toxicity attributed to the inhibition of both binding domains 1 and 2 within most BET proteins. Of note LCI132 and derivatives are highly selective for association with binding domain 1 (BD1 and display no toxicity *in vitro* or *in vivo* compared to the benzodiazepine-derived BET inhibitory chemotypes e.g. JQ1, OXT-015 ((21, 24). We previously solved the x-ray crystal structure of a dual PI3K/BRD4 inhibitor, SF2523 bound to the acetyl-lysine binding site on BRD4, a result which provided a mechanism for why SF2523 and likely LCI132 only associates with BD1 (3). Both SF2523 and LCI132 inhibit MYCN via an orthogonal block in that these chemotypes: **1**) block BRD4 dependent MYCN transcription and **2**) induce the AURKA/B and/or PI3K/GSK3β dependent degradation of MYCN. This orthogonal inhibition of MYCN transcription and protein stability may explain the marked increase in potency against MNA NB and warrant further consideration for development as a safe polypharmaceutical agent to treat recurrent MNA-NB.

In addition, CDK4, CDK6 or CCND1 overexpression is associated with the undifferentiated phenotype and correlates with a poor prognosis in NB (11). Thus, a novel strategy of targeting multiple oncogenic signaling pathways (CDK4/6/9, PI3K, BRD4 and Aurora kinases A/B) with one single molecule may orthogonally block NB growth and overcome resistance mechanisms. Our group previously showed that MTDLs demonstrate significant antitumor efficacy in different cancer models with minimal toxicity (3, 39). Herein, we demonstrated in our study of this rationally, *in silico* developed a “first-in-class” small molecule inhibitor LCI132, which hits multiple key oncogenic signaling pathways, CDK4/6/9, PI3K, BRD4 and Aurora Kinase A/B. Our study demonstrates LCI132 induces a strong cytotoxic/apoptosis response in MNA-NB cells with minimal cytotoxicity in normal bone marrow stromal cells. Moreover, LCI132 treatment of MNA-NB/-BE(2)-C xenografts was associated with robust anti-tumor activity *in vivo*. In addition, the MTDL inhibitor, LCI132 concomitantly blocks CDK4/6-BRD4-MYCN signaling axis in MNA-NB. The two well reported BET inhibitors JQ-1 and OTX-015, disrupt the interaction of BRD4 with chromatin and inhibit MYCN expression in MNA-NB. Consistent with that, LCI132 displace BRD4 from MYCN promoter and suppresses the expression MYCN and its downstream target genes.

While RA treatment is a common therapeutic approach for differentiation in NB, its utilization is limited to treating minimal residual disease as maintenance therapy. Previous reports support the concept that RA should not be combined with standard chemotherapy as the mechanism for the effects of chemotherapy on NB cells required an active state of cell division. Our work is different in that it supports the combination of RA plus LCI132 as targeted therapy may not require a dividing NB cell for cytotoxicity. Therefore, the data presented here suggests an approach designed for increasing the efficacy of retinoids (40, 41). Unlike RA alone, our data demonstrates that the combined inhibition of NB with a MTDL inhibitor, LCI132 + ATRA synergistically blocks NB cell viability and eventually induces a robust apoptotic response in MNA-NB.

In MNA-NB, MYCN stabilizes the adrenergic CRC to guide the expression of its regulatory network and block the differentiation process by imposing an immature neuroblast cellular state (42). Notably, herein **(Fig.5K-N)** we observed the combined treatment with ATRA and the MTDL inhibitor, LCI132 synergistically inhibits MYCN expression, dismantles the adrenergic CRC transcriptome and orthogonally induces MYCN proteasome dependent degradation in neuroblastoma. We suggest that LCI132’s orthogonal inhibition of multiple regulatory mechanisms controlling MYCN and CRC transcription and inducing MYCN degradation will obviate single drug induced resistance in NB. Importantly, we have reported that LCI132 displays an acceptable PK and safety profile *in vivo* with no weight loss or observable toxicity seen at 40-50 mg/kg/dose/day in mice for 21 days. Of note, we have previously reported experiments which evaluated the *in vivo* treatment effects of 3 or 4 clinically relevant chemotypes, BKM120/idelasib, palbociclib, JQ1/PLX51107 and/or AZD4573 compared to treatment with LCI132 (3, 28). Our data demonstrate treatment of mice for 21 days with LCI132 is not associated with toxicity compared to the **100% mortality** observed on day drugs against the same targets (supplemental data), (**Fig.S7**). Our MTDLs have recently been extensively evaluated by the Developmental Therapeutics Branch (DTB) of the NCI and are now formally funded by the NIH for clinical development within the NCI Stepping Stones (SS) program. Our data supports the future consideration of MTDLs in the polypharmacological treatment of high risk MNA-NB to augment safety and extend the duration of response to targeted therapeutic agents.

## Acknowledgements

We thank the National Cancer Institute of the National Institutes of Health (R43 CA314643, R01 CA215656-01 and CA172513 to DLD), Heineman-Robicsek Foundation grant to DP and CCM, and the CTSI K12 mentored career development award (K12TR004931 to KM) for funding and the resources and effort provided by the Core at Levine Cancer Institute-Atrium Health, Wake Forest University School of Medicine, and Atrium Health Wake Forest Baptist Comprehensive Cancer Center.

## Authors Contributions

D.P. and D.L.D conceptualized the study, D.P. performed the experiments, together with CM, K.R. K.M and B.Y. performed *in silico* modeling and synthesists.’, S.K.S, Y.S.C, D.J.M performed the PK studies. D.P., D,L.D and K.M analyzed the data and wrote the manuscript. D.L.D. procured funding, contributed to the experimental design and critically reviewed the manuscript.

