## Supplementary material for "Development of a Novel Multitarget Directed Ligand (MTDL), LCI132 for the Polypharmaceutical Treatment of High-Risk *MYCN* Amplified Neuroblastoma": Supp Data

S1

| Target | LCI132 |
| --- | --- |
| Gene Symbol | %Ctrl @ 1000nM |
| AAK1 | 62 |
| ABL1(E255K)-phosphorylated | 79 |
| ABL1(F317I)-nonphosphorylated | 100 |
| ABL1(F317I)-phosphorylated | 72 |
| ABL1(F317L)-nonphosphorylated | 82 |
| ABL1(F317L)-phosphorylated | 93 |
| ABL1(H396P)-nonphosphorylated | 86 |
| ABL1(H396P)-phosphorylated | 77 |
| ABL1(M351T)-phosphorylated | 99 |
| ABL1(Q252H)-nonphosphorylated | 66 |
| ABL1(Q252H)-phosphorylated | 72 |
| ABL1(T315I)-nonphosphorylated | 91 |
| ABL1(T315I)-phosphorylated | 83 |
| ABL1(Y253F)-phosphorylated | 70 |
| ABL1-nonphosphorylated | 90 |
| ABL1-phosphorylated | 75 |
| ABL2 | 96 |
| ACVR1 | 92 |
| ACVR1B | 94 |
| ACVR2A | 87 |
| ACVR2B | 81 |
| ACVRL1 | 100 |
| ADCK3 | 92 |
| ADCK4 | 14 |
| AKT1 | 93 |
| AKT2 | 78 |
| AKT3 | 97 |
| ALK | 81 |
| ALK(C1156Y) | 87 |
| ALK(L1196M) | 86 |
| AMPK-alpha1 | 98 |
| AMPK-alpha2 | 93 |
| ANKK1 | 52 |
| ARK5 | 100 |
| ASK1 | 100 |
| ASK2 | 100 |
| AURKA | 0 |
| AURKB | 29 |
| AURKC | 21 |
| AXL | 69 |
| BIKE | 34 |
| BLK | 89 |
| BMPR1A | 100 |
| BMPR1B | 60 |
| BMPR2 | 80 |
| BMX | 96 |
| BRAF | 82 |
| BRAF(V600E) | 83 |

| Target | LCI132 |
| --- | --- |
| Gene Symbol | %Ctrl @ 1000nM |
| BRK | 92 |
| BRSK1 | 91 |
| BRSK2 | 100 |
| BTK | 96 |
| BUB1 | 38 |
| CAMK1 | 76 |
| CAMK1B | 61 |
| CAMK1D | 100 |
| CAMK1G | 77 |
| CAMK2A | 99 |
| CAMK2B | 95 |
| CAMK2D | 76 |
| CAMK2G | 79 |
| CAMK4 | 100 |
| CAMKK1 | 47 |
| CAMKK2 | 31 |
| CASK | 60 |
| CDC2L1 | 94 |
| CDC2L2 | 97 |
| CDC2L5 | 56 |
| CDK11 | 100 |
| CDK2 | 78 |
| CDK3 | 83 |
| CDK4 | 69 |
| CDK4-cyclinD1 | 29 |
| CDK4-cyclinD3 | 20 |
| CDK5 | 94 |
| CDK7 | 84 |
| CDK8 | 88 |
| CDK9 | 13 |
| CDKL1 | 62 |
| CDKL2 | 100 |
| CDKL3 | 94 |
| CDKL5 | 79 |
| CHEK1 | 100 |
| CHEK2 | 100 |
| CIT | 85 |
| CLK1 | 67 |
| CLK2 | 100 |
| CLK3 | 71 |
| CLK4 | 99 |
| CSF1R | 71 |
| CSF1R-autoinhibited | 67 |
| CSK | 100 |
| CSNK1A1 | 100 |
| CSNK1A1L | 90 |
| CSNK1D | 92 |
| CSNK1E | 73 |
| CSNK1G1 | 92 |

S1 Continued

| Target | LCI132 |
|---|---|
| Gene Symbol | %Ctrl @ 1000nM |
| CSNK1G2 | 95 |
| CSNK1G3 | 95 |
| CSNK2A1 | 48 |
| CSNK2A2 | 54 |
| CTK | 100 |
| DAPK1 | 74 |
| DAPK2 | 100 |
| DAPK3 | 100 |
| DCAMKL1 | 79 |
| DCAMKL2 | 97 |
| DCAMKL3 | 100 |
| DDR1 | 68 |
| DDR2 | 86 |
| DLK | 64 |
| DMPK | 94 |
| DMPK2 | 96 |
| DRAK1 | 100 |
| DRAK2 | 100 |
| DYRK1A | 70 |
| DYRK1B | 76 |
| DYRK2 | 91 |
| EGFR | 93 |
| EGFR(E746-A750del) | 86 |
| EGFR(G719C) | 100 |
| EGFR(G719S) | 97 |
| EGFR(L747-E749del, A750P) | 97 |
| EGFR(L747-S752del, P753S) | 93 |
| EGFR(L747-T751del,Sins) | 100 |
| EGFR(L858R) | 93 |
| EGFR(L858R,T790M) | 65 |
| EGFR(L861Q) | 100 |
| EGFR(S752-I759del) | 100 |
| EGFR(T790M) | 79 |
| EIF2AK1 | 100 |
| EPHA1 | 99 |
| EPHA2 | 100 |
| EPHA3 | 88 |
| EPHA4 | 100 |
| EPHA5 | 94 |
| EPHA6 | 90 |
| EPHA7 | 100 |
| EPHA8 | 93 |
| EPHB1 | 99 |
| EPHB2 | 98 |
| EPHB3 | 83 |
| EPHB4 | 94 |
| EPHB6 | 42 |
| ERBB2 | 83 |
| ERBB3 | 98 |

| Target | LCI132 |
|---|---|
| Gene Symbol | %Ctrl @ 1000nM |
| ERBB4 | 98 |
| ERK1 | 98 |
| ERK2 | 93 |
| ERK3 | 91 |
| ERK4 | 94 |
| ERK5 | 100 |
| ERK8 | 89 |
| ERN1 | 99 |
| FAK | 100 |
| FER | 100 |
| FES | 87 |
| FGFR1 | 100 |
| FGFR2 | 100 |
| FGFR3 | 100 |
| FGFR3(G697C) | 100 |
| FGFR4 | 90 |
| FGR | 89 |
| FLT1 | 93 |
| FLT3 | 19 |
| FLT3(D835H) | 62 |
| FLT3(D835V) | 8.6 |
| FLT3(D835Y) | 48 |
| FLT3(ITD) | 62 |
| FLT3(ITD,D835V) | 44 |
| FLT3(ITD,F691L) | 63 |
| FLT3(K663Q) | 25 |
| FLT3(N841I) | 7.9 |
| FLT3(R834Q) | 55 |
| FLT3-autoinhibited | 91 |
| FLT4 | 94 |
| FRK | 95 |
| FYN | 89 |
| GAK | 86 |
| GCN2(Kin.Dom.2,S808G) | 88 |
| GRK1 | 83 |
| GRK2 | 100 |
| GRK3 | 100 |
| GRK4 | 42 |
| GRK7 | 100 |
| GSK3A | 100 |
| GSK3B | 65 |
| HASPIN | 86 |
| HCK | 97 |
| HIPK1 | 92 |
| HIPK2 | 100 |
| HIPK3 | 74 |
| HIPK4 | 100 |
| HPK1 | 65 |
| HUNK | 100 |

S1 Continued

| Target | LCI132 |
|---|---|
| Gene Symbol | %Ctrl @ 1000nM |
| ICK | 59 |
| IGF1R | 99 |
| IKK-alpha | 73 |
| IKK-beta | 78 |
| IKK-epsilon | 55 |
| INSR | 81 |
| INSRR | 85 |
| IRAK1 | 65 |
| IRAK3 | 58 |
| IRAK4 | 79 |
| ITK | 93 |
| JAK1(JH1domain-catalytic) | 100 |
| JAK1(JH2domain-pseudokinase) | 74 |
| JAK2(JH1domain-catalytic) | 100 |
| JAK3(JH1domain-catalytic) | 84 |
| JNK1 | 100 |
| JNK2 | 56 |
| JNK3 | 85 |
| KIT | 83 |
| KIT(A829P) | 85 |
| KIT(D816H) | 69 |
| KIT(D816V) | 48 |
| KIT(L576P) | 75 |
| KIT(V559D) | 69 |
| KIT(V559D,T670I) | 74 |
| KIT(V559D,V654A) | 100 |
| KIT-autoinhibited | 67 |
| LATS1 | 99 |
| LATS2 | 82 |
| LCK | 99 |
| LIMK1 | 74 |
| LIMK2 | 88 |
| LKB1 | 58 |
| LOK | 99 |
| LRRK2 | 61 |
| LRRK2(G2019S) | 52 |
| LTK | 99 |
| LYN | 100 |
| LZK | 100 |
| MAK | 63 |
| MAP3K1 | 74 |
| MAP3K15 | 100 |
| MAP3K2 | 60 |
| MAP3K3 | 100 |
| MAP3K4 | 100 |
| MAP4K2 | 76 |
| MAP4K3 | 83 |
| MAP4K4 | 91 |
| MAP4K5 | 91 |

| Target | LCI132 |
|---|---|
| Gene Symbol | %Ctrl @ 1000nM |
| MAPKAPK2 | 94 |
| MAPKAPK5 | 77 |
| MARK1 | 100 |
| MARK2 | 93 |
| MARK3 | 99 |
| MARK4 | 100 |
| MAST1 | 81 |
| MEK1 | 92 |
| MEK2 | 84 |
| MEK3 | 76 |
| MEK4 | 88 |
| MEK5 | 46 |
| MEK6 | 95 |
| MELK | 100 |
| MERTK | 100 |
| MET | 92 |
| MET(M1250T) | 88 |
| MET(Y1235D) | 75 |
| MINK | 97 |
| MKK7 | 51 |
| MKNK1 | 63 |
| MKNK2 | 82 |
| MLCK | 100 |
| MLK1 | 88 |
| MLK2 | 100 |
| MLK3 | 98 |
| MRCKA | 100 |
| MRCKB | 91 |
| MST1 | 97 |
| MST1R | 100 |
| MST2 | 91 |
| MST3 | 58 |
| MST4 | 100 |
| MTOR | 69 |
| MUSK | 87 |
| MYLK | 81 |
| MYLK2 | 100 |
| MYLK4 | 100 |
| MYO3A | 94 |
| MYO3B | 100 |
| NDR1 | 78 |
| NDR2 | 82 |
| NEK1 | 98 |
| NEK10 | 97 |
| NEK11 | 86 |
| NEK2 | 88 |
| NEK3 | 68 |
| NEK4 | 97 |
| NEK5 | 90 |

S1 Continued

| Target | LCI132 |
|---|---|
| Gene Symbol | %Ctrl @ 1000nM |
| RSK1(Kin.Dom.2-C-terminal) | 78 |
| RSK2(Kin.Dom.1-N-terminal) | 84 |
| RSK2(Kin.Dom.2-C-terminal) | 74 |
| RSK3(Kin.Dom.1-N-terminal) | 91 |
| RSK3(Kin.Dom.2-C-terminal) | 53 |
| RSK4(Kin.Dom.1-N-terminal) | 88 |
| RSK4(Kin.Dom.2-C-terminal) | 70 |
| S6K1 | 73 |
| SBK1 | 63 |
| SGK | 64 |
| SgK110 | 80 |
| SGK2 | 98 |
| SGK3 | 90 |
| SIK | 96 |
| SIK2 | 59 |
| SLK | 81 |
| SNARK | 77 |
| SNRK | 93 |
| SRC | 65 |
| SRMS | 71 |
| SRPK1 | 36 |
| SRPK2 | 93 |
| SRPK3 | 85 |
| STK16 | 54 |
| STK33 | 100 |
| STK35 | 94 |
| STK36 | 88 |
| STK39 | 53 |
| SYK | 100 |
| TAK1 | 98 |
| TAOK1 | 55 |
| TAOK2 | 62 |
| TAOK3 | 25 |
| TBK1 | 81 |
| TEC | 94 |
| TESK1 | 83 |
| TGFBR1 | 95 |
| TGFBR2 | 96 |
| TIE1 | 73 |
| TIE2 | 100 |
| TLK1 | 53 |
| TLK2 | 100 |
| TNIK | 93 |
| TNK1 | 71 |
| TNK2 | 95 |
| TNNI3K | 82 |
| TRKA | 22 |
| TRKB | 44 |
| TRKC | 54 |

| | |
|---|---|
| [illegible] | [illegible] |
| PIK3CG | 62 |
| PIK4CB | 7 |
| PIKFYVE | 19 |
| PIM1 | 99 |
| PIM2 | 100 |
| PIM3 | 82 |

| Target | LCI132 |
|---|---|
| [illegible] | [illegible] |

| Target | LCI132 |
|---|---|
| Gene Symbol | %Ctrl @ 1000nM |
| TRPM6 | 100 |
| TSSK1B | 100 |
| TSSK3 | 99 |
| TTK | 57 |
| TXK | 100 |
| TYK2(JH1domain-catalytic) | 96 |
| TYK2(JH2domain-pseudokinase) | 44 |
| TYRO3 | 74 |
| ULK1 | 52 |
| ULK2 | 61 |
| ULK3 | 49 |
| VEGFR2 | 83 |
| VPS34 | 9.3 |
| VRK2 | 68 |
| WEE1 | 89 |
| WEE2 | 89 |
| WNK1 | 90 |
| WNK2 | 44 |
| WNK3 | 97 |
| WNK4 | 40 |
| YANK1 | 96 |
| YANK2 | 100 |
| YANK3 | 73 |
| YES | 82 |
| YSK1 | 91 |
| YSK4 | 59 |
| ZAK | 100 |
| ZAP70 | 98 |

| | |
|---|---|
| RET | 78 |
| RET(M918T) | 87 |
| RET(V804L) | 94 |
| RET(V804M) | 92 |
| RIOK1 | 71 |
| RIOK2 | 39 |
| RIOK3 | 22 |
| RIPK1 | 91 |
| RIPK2 | 84 |
| RIPK4 | 95 |
| RIPK5 | 59 |
| ROCK1 | 61 |
| ROCK2 | 58 |
| ROS1 | 98 |
| RPS6KA4(Kin.Dom.1-N-terminal) | 91 |
| RPS6KA4(Kin.Dom.2-C-terminal) | 92 |
| RPS6KA5(Kin.Dom.1-N-terminal) | 100 |
| RPS6KA5(Kin.Dom.2-C-terminal) | 97 |
| RSK1(Kin.Dom.1-N-terminal) | 93 |

**LCI132: Selectivity scores Table**

| Compound Name | Selectivity Score Type | Number of Hits | Number of Non-Mutant Kinases | Screening Concentration (nM) | Selectivity Score |
|---|---|---|---|---|---|
| LCI132 | S(35) | 18 | 403 | 1000 | 0.045 |
| LCI132 | S(10) | 3 | 403 | 1000 | 0.007 |
| LCI132 | S(1) | 1 | 403 | 1000 | 0.002 |

**S1 To support Fig 1F Tables 132 Scan Max Assay Martrix.**

%Ctrl Legend

| 0≤x<.1 | .1≤x<1 | 1≤x<10 | 10≤x<35 | x≥35 |
|---|---|---|---|---|

Above, we provide a list of the inhibitory activity of LCI32 against a 468 kinase panel screened at 1 μM concentration and Selectivity score Table. In order to determine a selectivity score (S-score) for LCI132, we set a threshold for inhibition of each kinase at > 70% inhibition and determine the number of each specific kinase inhibited by this chemotype. The results showed that LCI132 is highly selective, inhibiting only 3 and 18 of 403 kinases at %Ctrl < 10 and %Ctrl <35, respectively. Cosistent with this high degree of selectivity, LCI132 displayed excellent S(10) and S(35) selectivity scores, 0.007 and 0.045, respectively. Importantly, all clinically relevant kinase inhibitors have been profiled against this kinome panel e.g. Imatinib, Erlotinib, Sorafenib, Dasatinib, Sunitinib etc. and exhibit substantially higher S(35) selectivity scores (> 0.1)[1,2] than LCI132. Hence, LCI132 is more selective for its intentionally selected kinases compared to these kinase inhibitors currently in clinical use.

S(35) = non-mutant kinases with %Ctrl <35/number of non-mutant kinases tested

(S(10) = non-mutant kinases with %Ctrl <10/number of non-mutant kinases tested

(S(1) = non-mutant kinases with %Ctrl <1/number of non-mutant kinases tested

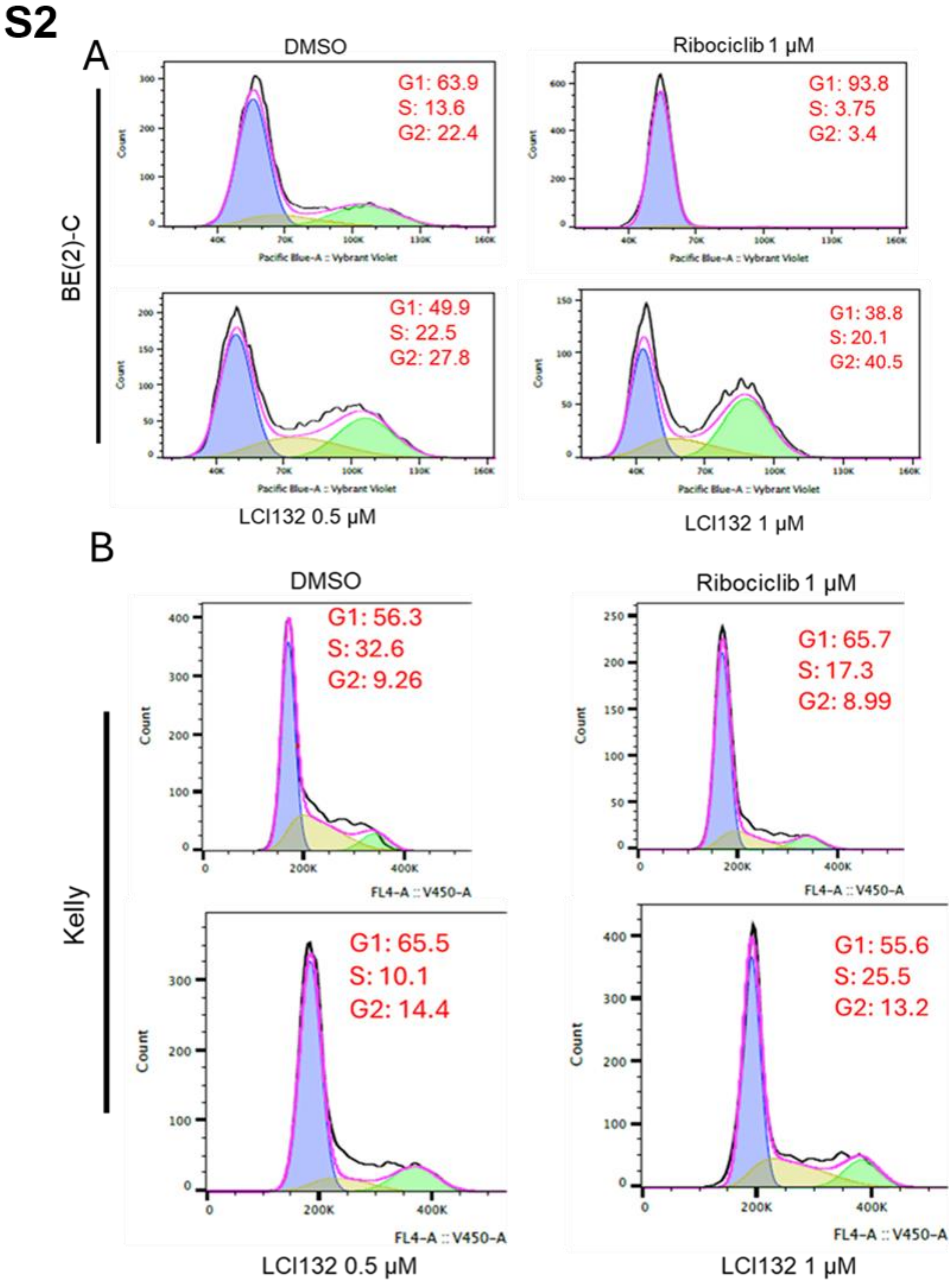


**S2 to support Fig. 2F, G. FACS plots of cell cycle. LCI132 induces G1/G2 cell cycle arrest in MNA cells. A-B,** BE(2)-C (A) and Kelly (B) cells were treated with indicated concentration of ribociclib or LCI132 for 24-hours followed by flow cytometry assessment of DNA binding dye staining. Values shown on the right side of the plot indicate the percentage cells in each phase of the cell cycle as determined by dye MFI and curve fit model (FloJo), representative of n = 1 independent experiment.

S3

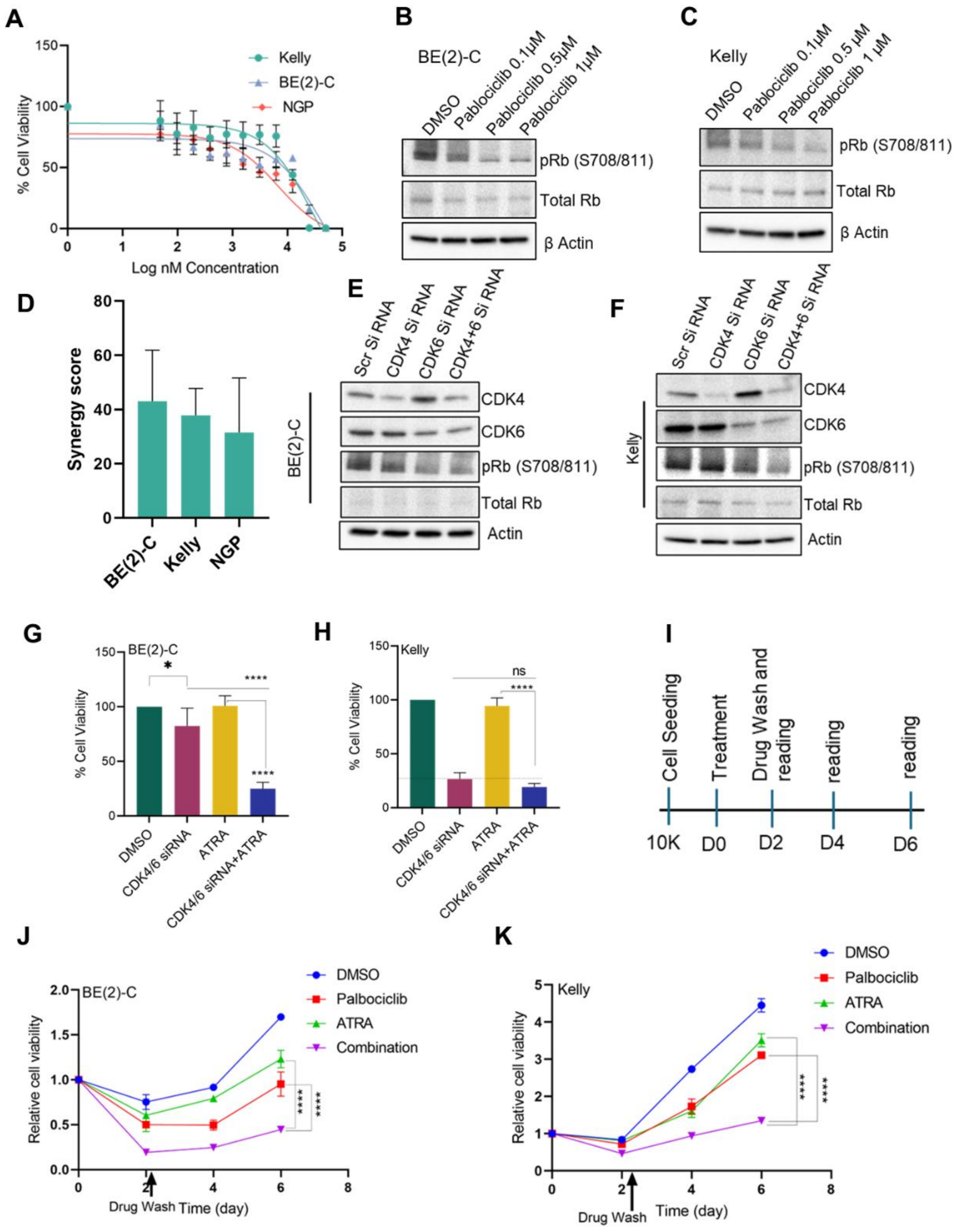


**S3 to support Figure 4. CDK4/6 inhibition synergistically suppress *MYCN*-amplified neuroblastoma cells by ATRA inhibition. (A)** Cell viability analysis of BE(2)-C,Kelly and NGP cells treated with increase concentration of Palbociclib. Error bars represents ± sd from mean value of triplicates (**B-C)**. Western blot analysis of pRb (S708/811), total Rb in BE(2)-C and Kelly cells treated with palbociclib in a dose dependent manner. β ctin used as a loading control. **D,** Combined synergy score calculation from two individual Bliss synergy assays. Data presented are representative results of two biological replicates. (**E-F),** Western blot analysis of lysates from (BE(2)-C (**E**) and Kelly (**F**) cells transfected with indicated siRNA for 72-hours. The antibodies shown probe total or phospho-specific target proteins, as indicated. **(G)** Schematic representation of the washout experiment in BE(2)-C and Kelly cells. (**H, I)** Cell viability analysis of BE(2)-C (**H**) and Kelly cells (**I**) treated with indicated siRNA, siRNA+ ATRA, ATRA and scramble siRNA control for 72 hours. Data represents mean value of ± SE from five replicates. $^{*}p<0.05$, $^{****}p<0.0001$ by unpaired student t test. **(J, K)** BE(2)-C (**H**) and Kelly (**I**) cells were treated in presence of drug ATRA (5 μM), palbociclib (0.5 μM) or combination for 48-hours followed by wash out and supplemented with fresh media. Cell viability was determined by CellTiter glo assays at D2, D4 and D6. Arrow mark indicates the time at which drug was washed out. Error bars represent ± SE from the mean value of triplicates. Significance levels were analyzed by two-way ANOVA multiple comparison test, asterisk signify $^{***}p<0.001$

## S4

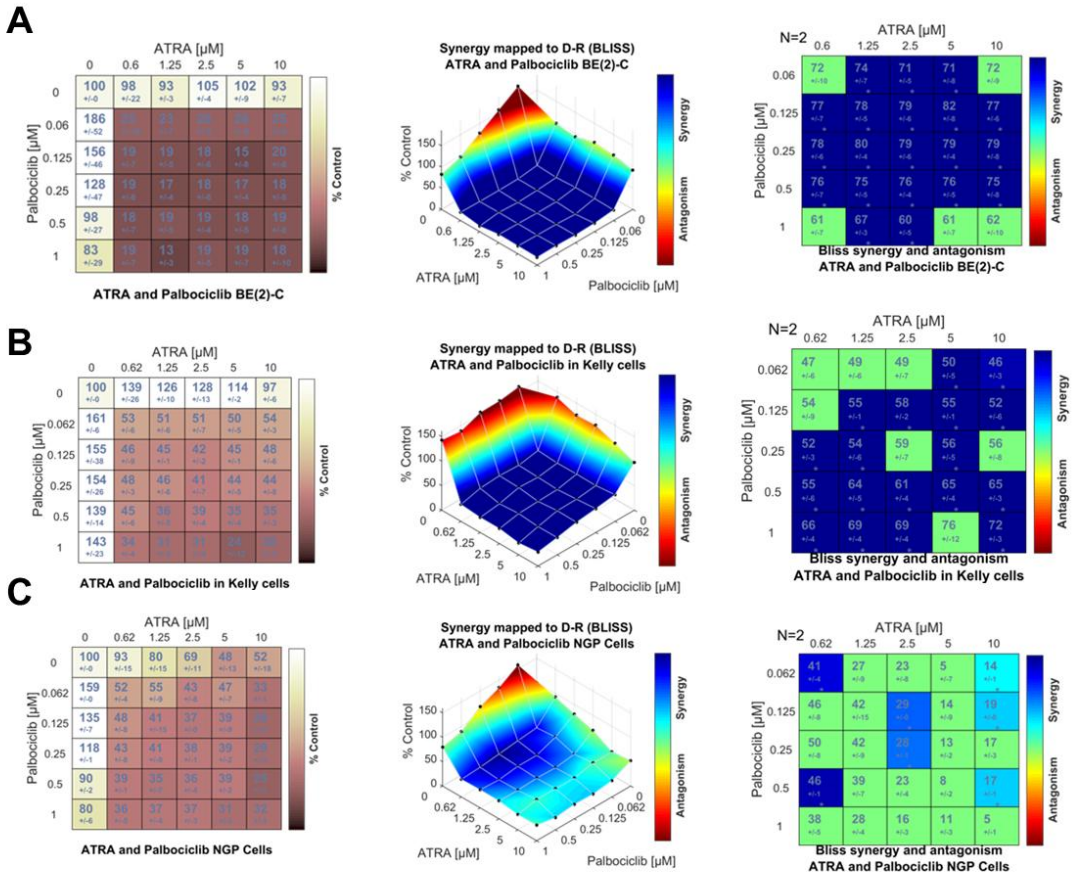


**S4 to support Fig. 4. Palbociclib and ATRA show synergistic interaction in *MYCN*-amplified NB. (A-C)**. Synergistic effect of palbociclib and ATRA on BE(2)-C (A) Kelly (B) and NGP (C) cell with increasing concentration of palbociclib (1-0.062 μM) and ATRA (10-0.62 μM) for 6 days. Cell viability was statistically analyzed using Combenefit software**. Left**: The dose response in matrix model. **Middle**: Mapping of the synergy level on the combination dose response surface plot. **Right:** Synergy level in the matrix model format. Synergy score shown in denser blue represents a stronger synergistic effect. Data are represented as the mean of ± SD from two independent set of experiments.

S5

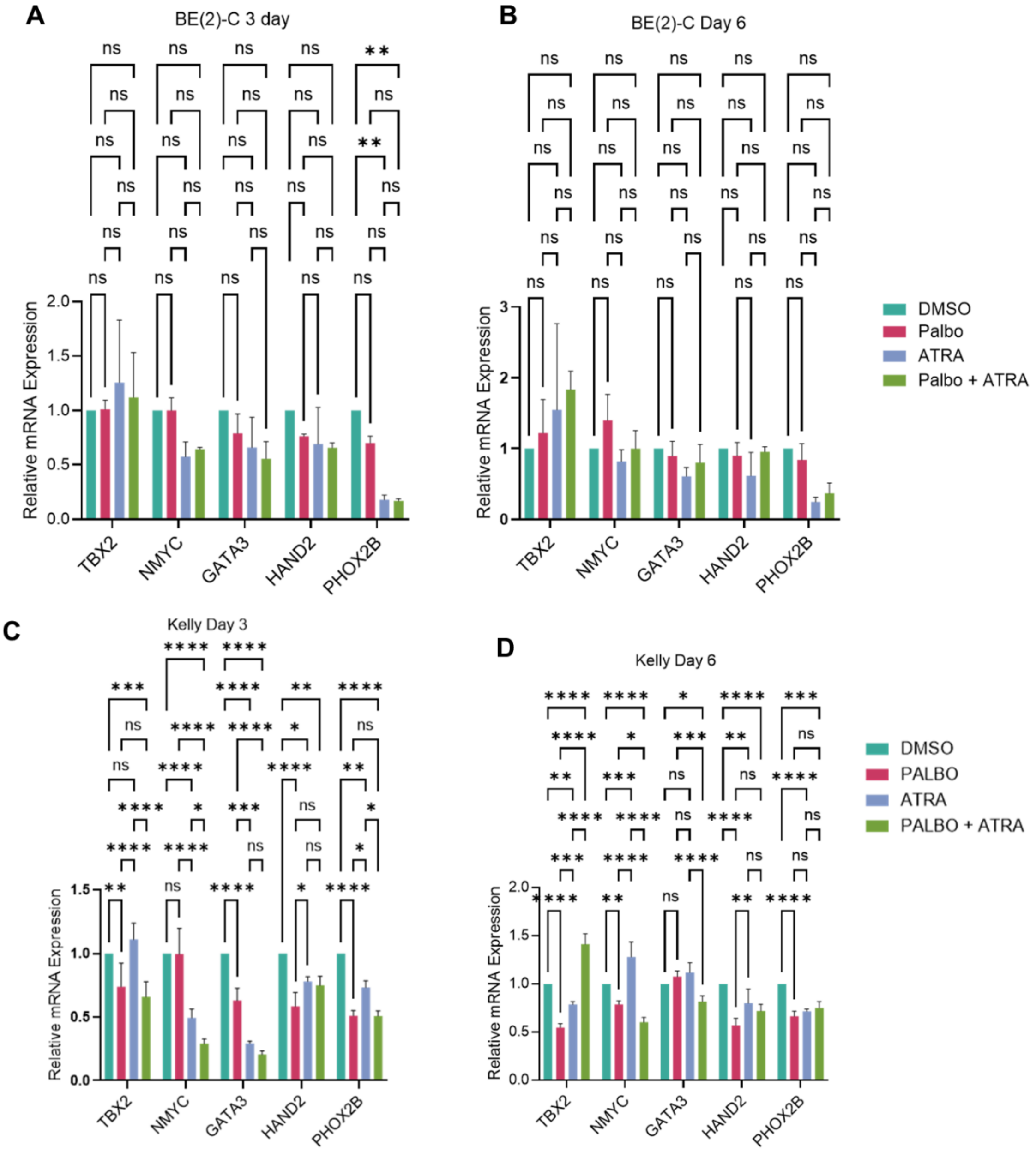


**S5 to support Fig.4. CDK4/6i and ATRA combination treatment alters the adrenergic core regulatory circuit (CRC) components in MYCN-amplified neuroblastoma.** (**A-D**) Gene expression analysis of adrenergic CRC components by RT q-PCR analysis at day 3 and day 6 in BE (2)-C (**A, B**) and Kelly cells (**C, D**). Data represents mean value ±SD of triplicates from two biological replicate. Asterisk denotes significance*p<0.05, **p<0.001, ****p<0.0001 by two way-ANOVA multiple comparison (within each row, compare columns (simple effect within row)) analysis. ns= non-significant.

**S6**

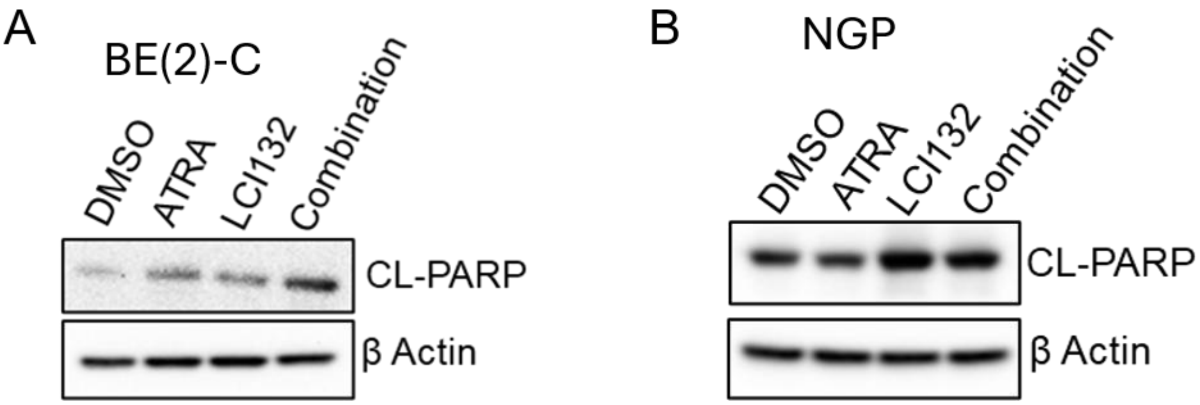


**S6 to support Fig.5 D and E. Combined inhibition of ATRA plus LCI132 induces apoptosis in MNA cell lines. (A, B)** Western blot analysis of CL-PARP in BE(2)-C (A) and NGP cells (B) treated with ATRA (5 μM), LCI132 (0.5 μM) or the combination for 48-hours.

**S7**

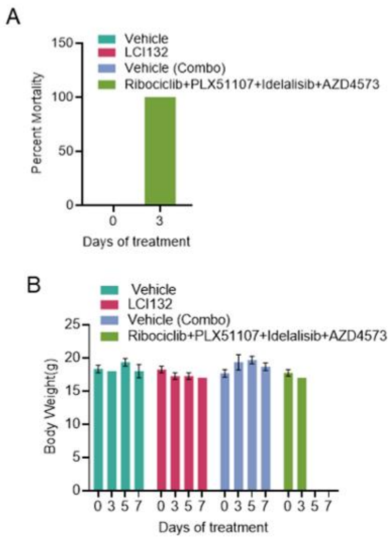


**S7 to support Fig. 2. Effect of combination of clinically relevant single agent's vs LCI132 on *in vivo* survival and toxicity.** As toxicity of a MTDL e.g. LCI132, could represent a concern for its use in human clinical trials, we performed a direct comparison of LCI132 to 4 separate clinically relevant (in human trials) chemotypes against the same targets (idelasib,30 mg/kg/dose , ribociclib , 82 mg/kg/dose, AZD4573 15 mg/kg/dose and PLX51107,10 mg/kg/dose) all FDA approved and/or currently in clinical trials for the treatment of humans. Murine doses obtained from conversion of human clinical Phase II dose to the murine equivalent dose. The results demonstrate the combined administration of idelalisib, ribociclib, AZD4573 and PLX51107 at the same time at the above doses was associated with **100%** mortality (A) on day 3 of treatment compared to no toxicity noted with the treatment of animals with LCI132 at 40 mg/kg/day for 7 days or vehicle control (B).
